# The future is fully defined: recombinant fragment E8 of laminin-511 is a viable xenofree alternative to Matrigel for hiPSC culture and differentiation into neurovascular cell types

**DOI:** 10.1101/2024.06.20.599891

**Authors:** Laís A. Ferreira, Denise Fabiano do Nascimento, Ishita Tandon, Lance Cordes, Kartik Balachandran

## Abstract

Matrigel remains the gold standard substrate for culture of induced pluripotent stem cells (iPSCs). However, its highly variable composition, animal origin and unpredictable effects on biological activity have been discussed for more than 3 decades. In this study, we explore the use of fragment E8 of recombinant laminin 511, commercially available in form of iMatrix-511, as an alternative to Matrigel for iPSC maintenance and differentiation. Female iMR90-4 human iPSCs were cultured on either iMatrix or Matrigel and assessed for cell growth and viability, pluripotency, genetic stability, and ability to differentiate into isogenic brain microvascular endothelial cells (iBMECs) and brain pericytes. It was observed that iMatrix facilitated iPSC growth and viability comparable to Matrigel while maintaining a higher number of more consistently sized colonies. Additionally, like Matrigel, iMatrix maintained the expression of pluripotency markers SSEA-4 and OCT-3/4 over 15 passages without inducing DNA damage. iMatrix also supported the differentiation of these iPSCs into isogenic iBMECs and pericytes, which were successfully co-culture for generation of a simplified blood-brain barrier model. Overall, we showed that iMatrix, which is a cost effective, fully defined, and xenofree alternative can be used as a substitute for Matrigel for maintenance and differentiation of iPSCs.

## Introduction

From layers of inactivated feeder cells [1] to humanized recombinant fragments of extracellular matrix (ECM) proteins with enhanced adhesion properties [2], the optimization of products and protocols for maintenance of induced pluripotent stem cells (iPSCs) in vitro is undeniably remarkable. With the constantly increasing interest in the study of iPSCs for therapeutic applications and disease modeling, various adhesion molecules and nutrient media compositions have been developed and validated, some in fully defined and xenofree formulations, contributing to overall better experimental reproducibility and robustness [3]. Currently, Matrigel (MG) is the most widely used ECM for iPSC culture, evidencing its broad applicability. However, this laminin-rich basement membrane preparation is secreted by mouse sarcoma cells, and therefore highly variable in composition and animal derived. In fact, the need to cautiously interpret data obtained from cells cultured on MG due to the biological activity of its undefined and variable components have been discussed since 1992 [4,5].

Considering that laminins, which are the major component of MG, play an essential role in the embryonic stem cell niche in vivo [6,7], the compatibility of several isoforms of laminin with pluripotent stem cell culture have been investigated based on the integrins expressed by these cells. In 2008, Miyazaki et al., pointed to the potential of human recombinant laminin-511 (LN511) as a suitable substrate for undifferentiated expansion of human embryonic stem cells, which was further confirmed by Rodin et al., 2010 and shown to promote superior cell adhesion than MG and other laminins while maintaining pluripotency and normal karyotypes [7,8]. A few years later, Miyazaki’s research on recombinant laminins led to the isolation of fragment E8 of LN511 (LN511-E8), the active integrin-binding site, and demonstration of its superior affinity to human pluripotent stem cells compared to its full-length counterpart [2].

Now, LN511-E8 is commercially available as iMatrix-511 (iM), a fully defined and xenofree product for efficient maintenance of pluripotent stem cells in vitro. In addition to the enhanced reproducibility resulting from the replacement of a poorly defined ECM mixture with a purified truncated recombinant protein, iM features other important advantages to known MG limitations, such as stability at room temperature, ease of manipulation, and improved cost effectiveness [4]. Another major benefit of iM adoption for stem cell research compared with other available products is that it allows for incorporation to the cell suspension immediately before seeding, making the laborious and time-consuming pre-coating step expendable and cell culture protocols substantially more flexible [9].

In this study, we aimed to investigate whether iM could be used as an alternative to MG in a set of previously validated protocols for maintenance and differentiation of human iPSCs into relevant cell types for blood-brain barrier (BBB) modeling. As hypothesized, based on previous research and the fact that laminins account for approximately 60% of MG composition [4], our results demonstrate that LN511-E8 efficiently supports iPSC culture while maintaining cell growth, pluripotency, and genetic stability comparable to their counterparts cultured on MG. As we delved deeper into the potential of these cells for tissue engineering applications, we have also demonstrated that iM could be successfully adopted in protocols validated using MG for directed differentiation into isogenic brain microvascular endothelial cells and brain pericytes. Overall, this study offers valuable insights into the potential adaptability of protocols for pluripotent stem cells research to increase experimental planning flexibility and facilitate compatibility with xenofree and fully defined approaches.

## Materials and methods

Details on cell culture reagents and primary antibodies used were included on S1 and S2 Tables, respectively.

### iPSC culture and coating strategies

A diagram illustrating the experimental design adopted in this study can be found in Fig 1. The cell line used was the female iMR90-4, purchased from WiCell. iPSCs were fed mTeSR+ media daily and passaged when approximately 80% confluency was observed. Upon first seeding after thawing, 10 mm Y-27632 (ROCK inhibitor) was added to the culture media – which was removed 24 hours later. For comparative assessment of coating strategies one vial of cryopreserved cells was split into iM and growth factor reduced MG cultures and carried out in parallel, as shown in Fig 1. These cultures will be referred to as iM-iPSCs and MG-iPSCs, respectively. Three pairs of cultures were used in independent experiments for collection of the presented data. iM was used at 0.25 µg/cm^2^ and added directly to the cell suspension immediately before seeding [9], while MG plates were pre-coated at 8.7 µg/cm^2^ [10] in DMEM/F-12 media at 37°C at least overnight and up to 10 days before cell seeding. Both cultures were passaged using ReLeSR for maintenance and seeded on 25 mm Thermanox coverslips (Electron Microscopy Sciences) every odd passage, from P35 to P49, for identification of pluripotency markers octamer binding transcription factor- 3/4 (OCT-3/4) [11] and the ganglioside stage-specific embryonic antigen-4 (SSEA-4) [12] through ICC techniques as described below.

**Fig 1.**
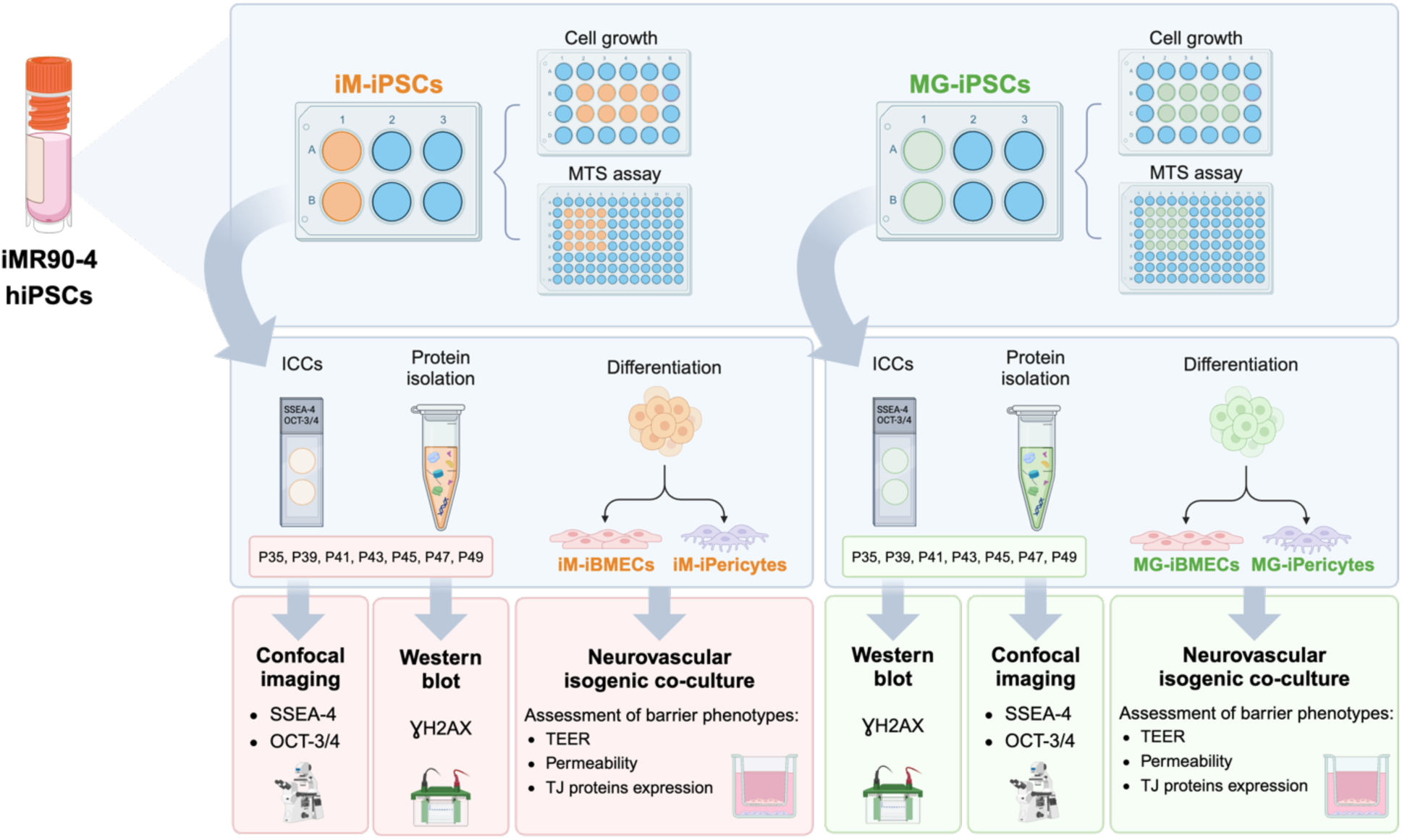
Overview of the experimental design adopted in the study. Three pairs of cultures derived from the same initial cryovial were carried out in parallel using iMatrix or Matrigel. Undifferentiated iPSCs in passages between 37 and 38 were used for comparative assessment of cell growth and viability through daily cell counts and MTS assays, respectively. In every odd passage between 35 and 49, iPSCs were seeded on Thermanox coverslips for the identification of expression of the pluripotency markers SSEA-4 and OCT-3/4 through ICC techniques, and protein samples were collected for assessment of γH2AX expression as an indicator of genetic stability. iPSCs grown using iM and MG were differentiated into iBMECs and iPericytes using the equivalent ECM when appropriate and used to establish neurovascular co-cultures of cells generated using each strategy. Finally, the development of barrier phenotypes was comparatively assessed through TEER, permeability to a small molecule, and expression of the tight junction proteins claudin-5 and zonula occludens-1 (ZO-1). iBMECs: iPSC-derived brain microvascular endothelial cells; ICC: immunocytochemistry; iM: iMatrix; MG: Matrigel; OCT-3/4: octamer binding transcription factor-3/4; P: passage; SSEA-4: stage-specific embryonic antigen-4; TEER: transendothelial electrical resistance; TJ: tight junction. Created with BioRender.

### Cell growth and viability assessment

For the analysis of cell growth, cells between P37 and P38 were lifted using ReLeSR and seeded onto 24-well plates at a density of 2x10^4^ cells/cm^2^ [9] in mTeSR+ following the coating strategies described above. In each of the following four days, the typical interval between passages, cells from two wells were imaged using an EVOS™ XL Core Imaging System, collected following 6 min of incubation with accutase and counted using an automated cell counter (Countess 3 FL, Thermo Fisher). Trypan blue was used to determine the number of live and dead cells at each evaluated timepoint. Pictures taken 24h post seeding were used for colony attachment analysis. Colonies were manually counted and measured using ImageJ (National Institutes of Health, Bethesda, Maryland, USA). All colonies with an area greater than or equal to 1500 µm² were considered for colony size assessment. Data was collected from nine images per group obtained in three independent experiments and reported as the average colony number and size per image analyzed.

For further quantification of cell viability, MTS assays were also performed using cultures between P37 and P38 according to the manufacturer instructions. In this case, 6.2x10^4^ cells/cm^2^ were seeded onto 96-well plates [13] in quadruplicates for each period in culture to be assessed. On the set timepoints, 20 µL of MTS was added to the wells containing 200 µL of cell culture media. After 3h of incubation, media was transferred to a new 96-well plate for absorbance reading at 490 nm. For every reading, wells containing only mTeSR+ and MTS (no cells) were also included for normalization of the obtained values. In both assays media in the remaining wells was changed daily until each experimental endpoint.

### iPSC derived brain microvascular endothelial cells (iBMECs)

Differentiations into neurovascular cell types were started in parallel using iPSCs grown on iM or MG and carried out using the same ECM in which initial undifferentiated cells were previously cultured as shown in Fig 1. The differentiation of iPSCs into iBMECs was performed based on a previously published method [14] summarized in Fig 5A and carried out in parallel using iM (iM-iBMECs) or MG (MG-iBMECs) as described above. Briefly, on day 0, iPSCs were incubated with accutase for 6 minutes at 37°C for singularization. Cells in suspension were collected, sampled for counting, and centrifuged at 300g for 5 minutes. The supernatant was then discarded, and the pellet was resuspended in mTeSR+ media containing 10 µM of ROCK inhibitor to generate a cell suspension at a density of 1.2x10^5^ cells/mL. Cells were either seeded onto MG pre-coated 6-well plates or uncoated plates after addition of iM to the suspension at a final density of 12.5x10^3^ cells/cm^2^.

On day 1 cells were given Essential 6 media (E6) to induce iBMEC differentiation, and daily media changes were performed until day 4. From day 5 to 7, cells received endothelial cell media (EC media), which consisted of human endothelial serum free medium supplemented with B27, containing 20 ng/mL of FGF-2 and 10 µM of retinoic acid with no media change on day 6 – characterizing the expansion phase. For further experimental use iBMECs were seeded onto 12 mm transwell inserts (Corning) and 15 mm Thermanox coverslips pre-coated with 0.4 mg/mL of collagen IV and 0.1 mg/mL of fibronectin, which were prepared on day 6 to allow for overnight incubation at 37°C. On day 7 the coating solution was aspirated, and plates were air dried for approximately 20 min before cell seeding.

For subculturing, cells were incubated with accutase for 1h at 37°C to promote cell detachment and singularization. Once in suspension, cells were collected, centrifuged in the above-mentioned conditions, and resuspended in EC media containing 10 µM retinoic acid. iBMEC were seeded based on a 1:3 split ratio, resulting in a density of approximately 2.5x10^6^ cells/transwell or coverslip. After 24 hours, all media was replaced with EC media without retinoic acid. The success of each differentiation process was assessed through the observation of expression of the tight junction protein zonula occludens-1 (ZO-1) through ICC techniques and daily transendothelial electrical resistance (TEER) measurements of cells seeded on coverslips and transwell inserts, respectively, as described below.

### iPSC derived brain pericytes (iPericytes)

Our protocol for the differentiation of iPSCs into brain pericytes (iPericytes), represented on Fig 6A, was developed based on a previously published two-step method in which iPSCs are initially differentiated into neural crest stem cells (iNCSC) before further differentiation into iPericytes [15]. For that, on day 0 iPSCs were singularized and seeded as described above for iBMEC differentiation, the only difference being a higher cell density of 9.1x10^4^ cells/cm^2^. iNCSC differentiation was induced on day 1, when media was switched to E6 supplemented with 1 µM CHIR99021, 10 µM SB431542, 0.01 µg/mL FGF-2, 1 µM dorsomorphin, and 22.5 µg/mL heparin – which will be referred to as E6-CSFD media. Cells were fed fresh E6-CSFD daily until day 15 and expanded in a 1:6 split ratio on days 3, 6, and 10 using accutase as previously described. ROCK inhibitor was added to the cultures at a final concentration of 10 µM for 24 hours following passaging on days 3 and 10 to increase cell attachment and viability at these critical time points of the differentiation process. Until day 15, cells were cultured on MG pre-coated plates or had iM added to cell suspensions immediately before seeding.

The first stage of the differentiation process was completed on day 15, when iNCSC cultures were purified through magnetic cell sorting based on the expression of nerve growth factor receptor (NGFR) according to manufacturer’s instructions. After sorting, iNCSC were seeded onto uncoated T-75 cell culture flasks (VWR) at a density of 1.4x10^4^ cells/cm^2^. Thermanox coverslips containing non-sorted iNCSCs were fixed with 4% paraformaldehyde (PFA; Electron Microscopy Sciences) and NGFR expression was demonstrated through ICCs. Between days 16 and 25, cells were fed fresh E6 supplemented with 10% fetal bovine serum (FBS) daily, and one passage was performed for expansion when cultures reached 90% confluency. For phenotype confirmation, on day 25 fully differentiated iPericytes were fixed and the expression of cell specific markers neural/glial antigen 2 (NG-2) and platelet-derived growth factor receptor β (PDGFR-β) was assessed through ICCs.

### Isogenic neurovascular co-culture

To comparatively assess the development of barrier phenotypes in neurovascular co-cultures of iPSC-derived cells generated using iM or MG, on experimental day 0 iBMECs and iPericytes differentiated on the same ECM were seeded onto the apical and basolateral chambers of 12 mm transwells kept on 12 well-plates as shown in Fig 7A. A diagram summarizing the timeline adopted for co-culture experiments can be found in Fig 7B. iPericytes were lifted from 75 cm^2^ cell culture flasks using accutase and processed following the steps described above to be seeded on the bottom of uncoated co-culture plates in E6 + 10% FBS at a density of 4.3x10^4^ cells/cm^2^. Day 7 iBMECs were subcultured onto collagen/fibronectin-coated transwell inserts in EC media + retinoic acid as previously detailed, which were then placed on iPericytes-seeded wells. On day 1, media in all samples was refreshed and retinoic acid was removed from iBMECs cultures. Thermanox coverslips, prepared according to cell type, were also included in every experiment and cells were seeded separately at equivalent cell densities for confirmation of iBMECs’ and iPericytes’ phenotypes through ICCs targeting ZO-1 and NG-2/PDGFR-β, respectively, as described below.

### Barrier tightness assessment

The development of barrier phenotypes was measured via TEER and permeability to 3 kDa dextran following the timeline shown in Fig 7B. An EVOM2 device (World Precision Instruments) was used to measure TEER in iBMECs monocultures and co-cultures with iPericytes over 5 days. Resistance was normalized through the subtraction of “blank” values collected from collagen/fibronectin-coated transwells containing culture media only before cells were seeded on day 0. For permeability assessment, 3kDa cascade blue-conjugated Dextran was diluted in EC media at a final concentration of 100 µg/ml and added to apical chambers (Fig 7A) of co-cultures on day 1. From day 2 to 5, 200 µl samples of E6 + 10% FBS were collected from the basolateral chamber and replaced with the same volume of fresh media. Samples were stored at -20°C until fluorescence intensity was measured using a Biotek Synergy Mix microplate reader at 400 ± 20 nm/450 ± 20 nm excitation/emission for determination of dextran concentrations in each sample.

For the establishment of a standard curve, dextran concentrations and the corresponding fluorescence intensities were analyzed. Technical duplicates were averaged and normalized by the subtraction of the baseline fluorescence intensity of the cell culture media in which samples were collected. Values were graphed to find the standard fit curves using MATLAB 2023a with the curve fitting toolbox add on. Using the fit function, the relationship between dextran concentrations and fluorescence intensities were calculated using the custom equation *y* = *a*(*x*^2^) + (*b* × *x*) with *y* being the dextran concentration, *x* the fluorescence intensity, and *a* and *b* values as the constants specific to each assay. After dextran concentrations were calculated, the relationship between dextran concentration and the time points of each sample collection were graphed. The slope 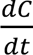 was calculated from the graph using the slope equation 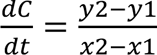 and adding a (0,0) point based on the assumption that the dextran concentration was 0 at time 0. The obtained values were then used to calculate the apparent permeability (Papp) through the following equation, where *Papp* is the apparent permeability (cm/sec), *Vr* the sample volume collected (ml), *A* the surface area of the porous membrane (cm^2^), *Co* the initial concentration of dextran in the EC media (mg/ml), and 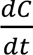 the slope of the dextran concentration over time (mg/(ml×sec)).

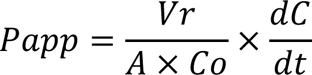

### Immunocytochemistry (ICC)

Cells to be analyzed via ICC techniques for immunoidentification of targets of interest were seeded on Thermanox coverslips – which were prepared following the same protocol used for the maintenance of each cell type in culture. iPSC cultures were fixed with 4% PFA for 10 min, while for iPSC-derived cells, PFA was added to the culture media at a 1% dilution and cells were incubated for 2 min before exposure to 4% PFA for 8 min. Following fixation, iPSCs and iPSC-derived cells were washed 3x with Phosphate-Buffered Saline (PBS) and Hank’s Balanced Salt Solution (HBSS), respectively, for 5 min under gentle orbital agitation. Blocking was performed at room temperature for 1h using a PBS-based blocking buffer composed of 0.1% gelatin from cold water fish skin (Sigma-Aldrich), 1% BSA, 2% normal goat serum (Thermo Fisher), 0.1% Triton X-100 (Millipore-Sigma), and 0.05% Tween 20 (Amresco) [16,17]. Serum, Triton X-100, and Tween-20 were not included in the blocking buffer for all markers and details on this step can be found on S2 Table, as well as all primary antibodies used and respective dilutions. All primaries were prepared in a solution of 1% BSA + 0.1% gelatin in PBS and allowed to bind to cellular epitopes overnight at 4°C. AlexaFluor (ThermoFisher) goat anti-mouse and anti-rabbit 488 (green fluorescence) and 594 (red fluorescence) were used as reporters at 1:100 and prepared in 1% BSA with 1:200 DAPI (Thermo Fisher) for nuclear staining. Following 1h of incubation at room temperature, coverslips were washed and mounted using ProLong Gold Antifade Mountant between a 75x25x1 mm microscope slide (VWR) and a 50x24x0.16 mm cover glass (Fisher Scientific) for image acquisition. Cells were imaged using an Olympus IX83 Inverted Confocal Microscope and all obtained files were further processed using ImageJ for conversion into image files and incorporation of scale bars.

### Western blot

For collection of protein samples, iPSCs and iBMECs were washed 3x with cold PBS or HBSS, respectively, for 5 min under gentle orbital agitation. Cells were lysed mechanically using cell scrapers while exposed to RIPA buffer containing a cocktail of protease inhibitors, sodium orthovanadate, and PMSF. Samples were then collected and centrifuged at 13,000 rpm for 10 min at 4°C for isolation of supernatants and removal of cell debris. Total protein content in cell lysates was determined using the Pierce Micro BCA Protein Assay Kit for normalization of protein loading concentration for electrophoresis. Samples collected from undifferentiated iPSCs every odd passage from 35 to 49 were loaded onto 4–20% Criterion TGX Precast Protein Gels (Bio-Rad) at a total protein concentration of 50 µg. SDS-PAGE was performed at 200V for 50 min at 4°C on ice. Proteins were then transferred to 0.45 µm pore size Immobilon-FL PVDF membranes (Millipore-Sigma) using a wet tank system at 10V for 16h at 4°C that included a cooling pack and a magnetic stir plate under constant agitation to keep the system from heating up. Membranes were stained with Ponceau S (Sigma-Aldrich) for 10 min to allow for visualization of proteins and assessment of transfer success. Blocking was carried out for 1h using Intercept (PBS) Protein-Free Blocking Buffer (Li-Cor) before membranes were incubated with anti-γH2AX overnight at 4°C for the assessment of DNA damage accumulation over 15 passages in culture.

Protein samples were also isolated from iBMECs grown on transwell inserts in co-culture with iPericytes following the steps described above and 40 µg of total protein extract were separated by molecular weight using 10% Criterion TGX Precast Protein Gels (Bio-Rad). In this case transfers were carried out at 80V for 1.5h and membranes were also blocked for 1h before incubation with primary antibodies targeting tight junction proteins claudin-5 and ZO-1. Blots were imaged using an Odyssey Imager (Li-Cor) following labeling with fluorophore-conjugated IRDye® 700CW and 800CW (Li-Cor) secondary antibodies. Densitometric analysis for protein expression quantification was performed using ImageJ and GAPDH densitometry values were used for signal normalization. For determination of the lane normalization factor, the value corresponding to the signal intensity of GAPDH for each sample was divided by the highest GAPDH signal on the blot. Data is presented as relative target protein signal, which was calculated by dividing the signal intensity by the lane normalization factor.

### Statistical analysis

All data sets were obtained from three experiments performed independently. GraphPad Prism version 10.1.1 for Mac OS (GraphPad Software, Boston, Massachusetts USA) was used to assess statistical significance and plot all graphs for data visualization. Data distribution was analyzed through Shapiro-Wilk and proper statistical tests were defined accordingly. Details on sample sizes and statistical tests corresponding to each data set are described on Fig captions. Differences were considered significant if p ≤ 0.05.

## Results

### iM facilitates the formation of a higher number of more consistent sized colonies of iPSCs

For a comparative assessment of colony attachment patterns and cell proliferation in iPSCs grown using iM and MG, cells were seeded onto 24-well plates at a density of 2x10^4^ cells/cm^2^ and in the following four days, the typical interval between passages for maintenance [3], phase contrast images were obtained as well as cell counts. Analysis of images acquired 24h after seeding (Fig 2A) showed that iM-iPSCs formed 42.75% more colonies than MG-iPSCs (p=0.0029), as cells seeded using iM formed 127.8 ± 11.4 colonies compared to 82.8 ± 5.9 formed by their MG-grown counterparts. When it comes to sizes, a trend towards larger colonies in MG-iPSCs was observed, however with no significance (p=0.0907). Interestingly, when comparing the standard error of the mean, the F test showed that colony sizes were more variable for MG-iPSCs than iM-iPSCs (p=0.0265).

**Fig 2.**
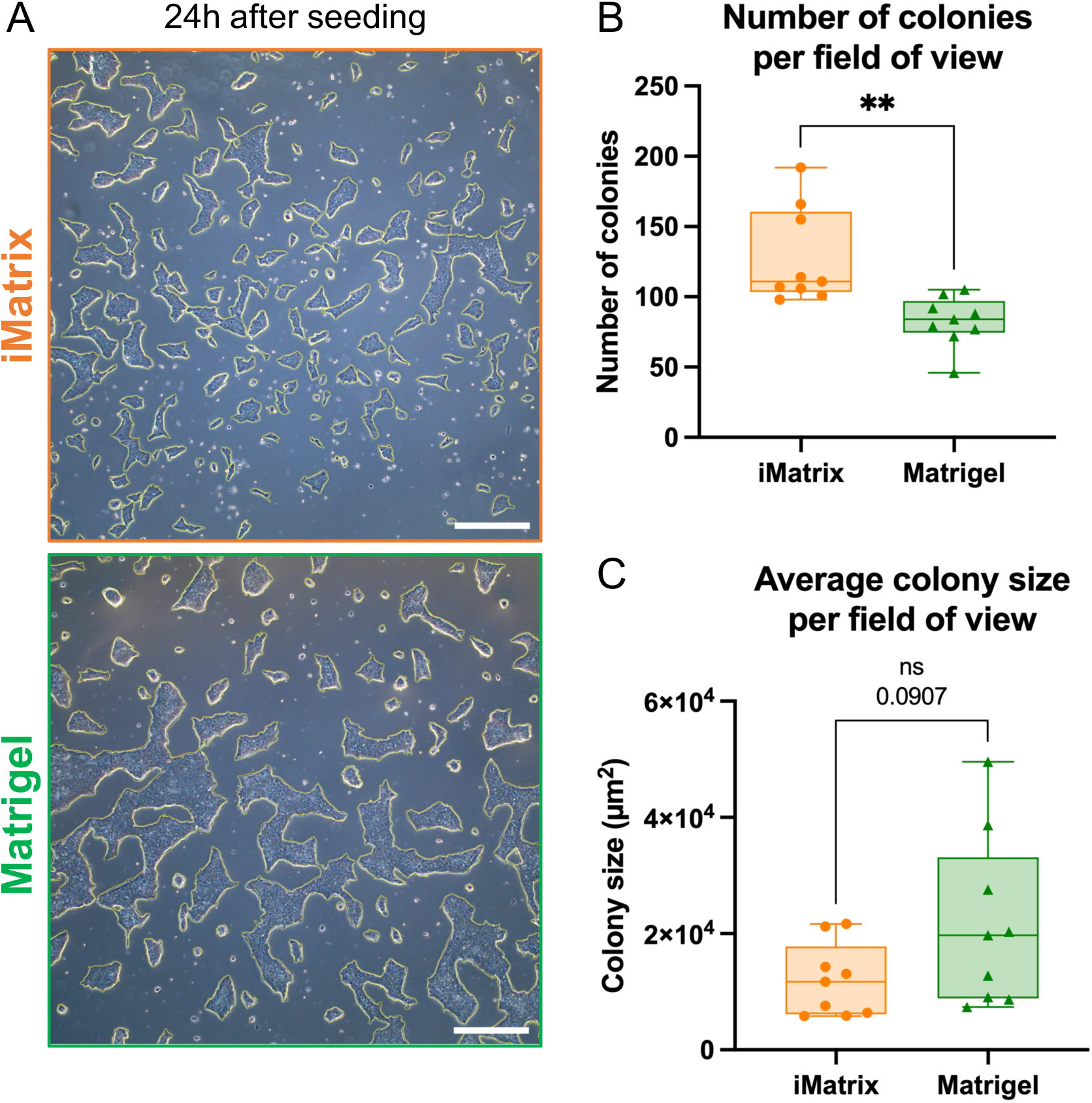
Colony formation analyses. Colony attachment was analyzed using **(A)** phase contrast microscopy images taken 24h after cell seeding. Colonies were counted and outlined for determination of surface area using ImageJ. The **(B)** number and **(C)** size of colonies are represented as Box & Whisker plots where the lines range from minimum to maximum values and the boxes depict the lower to upper quartile. Data was collected from 9 fields of view imaged in 3 independent experiments for each group and statistical significance was assessed through unpaired t tests for both datasets. **p=0.0029. ns: not significant.

The growth of iPSC colonies on both ECMs over four days can be visualized in the representative phase contrast images included on Fig 3A. In each timepoint wells were gently washed with D-PBS and the adhered cells were suspended and singularized using accutase for quantification using an automated cell counter. The progression in total cell number for each group over the assessed time in culture is shown in Fig 3B. Two-way ANOVA was adopted to accommodate all relevant variables, which showed that days in culture was the only significant source of variation for the evaluated data (p<0.0001), accounting for 69.3% of variation. The ECM used was associated with 0.53% of variation (p=0.3995) and the interaction between both factors for 0.37% (p=0.9184). Tukey’s multiple comparisons post-hoc test was also performed and further confirmed that the total cell number was statistically equivalent for iM-iPSCs and MG-iPSCs in all timepoints assessed, although averaged counts were higher for iM-iPSCs from day 1 to 4. In addition, both groups increased significantly in cell number over 4 days in culture in an equivalent manner. On day 4, the number of iM-iPSCs and MG-iPSCs increased 10.7x (p<0.0001) and 11.3x (p<0.0001), respectively, compared to data collected on day 1 – ranging from 94,490 ± 25,385 to 1,009,000 ± 150,811 for iM and 79,530 ± 22,538 to 899,333 ± 167,889 for MG.

**Fig 3.**
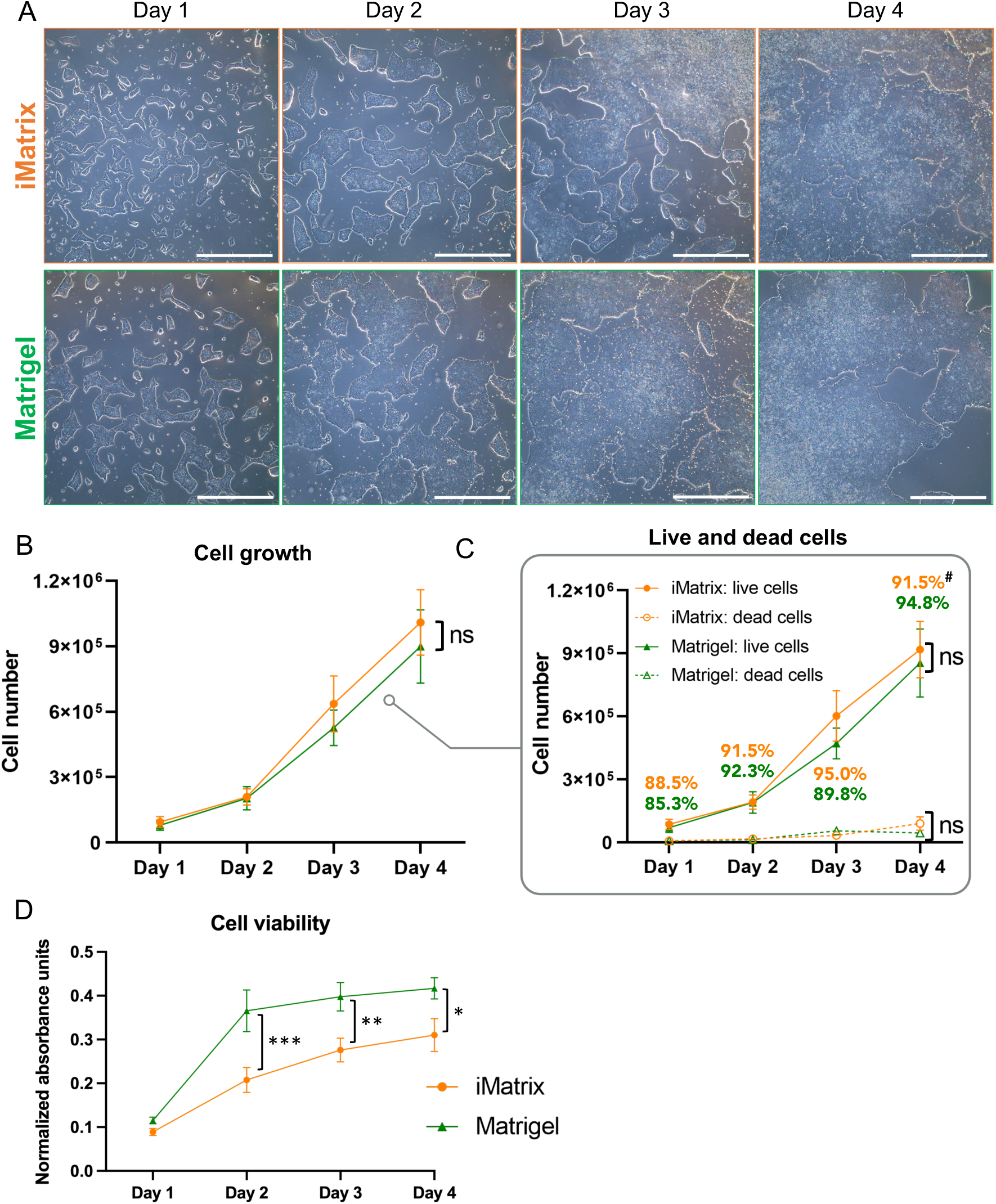
Growth and viability of iPSC cultures grown using iMatrix and Matrigel. **(A)** Representative phase contrast images of cells from both groups over four days in culture. Scale bars: 500 mm. The data from cell counts performed daily is shown as **(B)** total cell counts and **(C)** the number of live and dead cells determined by an automated cell counter based on trypan blue staining (n=6 counts/timepoint performed in 3 independent experiments). **#**percentage of live cells in each timepoint for cells grown using iMatrix (top, orange) or Matrigel (bottom, green). Cell viability was further assessed through **(D)** MTS assays (n=12 readings/timepoint performed in 3 independent experiments. All data was analyzed through two-way ANOVAs followed by Tukey’s multiple comparisons and is represented as mean ± SEM. *p=0.0125; **p=0.0045; ***p=0.0003. ns: not significant.

### iM supported similar increase in cell number to MG

With the incorporation of trypan blue in the cell counts described above, the number of live and dead cells in each time point was also determined. As shown in Fig 3C, the number of both live and dead cells was statistically equivalent for iM- and MG-iPSCs across all assessed timepoints. Within each ECM, the number of dead cells remained constant for four days, while the population of live cells progressively grew. The number of live cells increased significantly from days 2 to 3 (p=0.0002 for iM and p=0.0192 for MG) and 3 to 4 (p=0.0066 for iM and p=0.0006 for MG) in both groups, which cumulatively led to a 10.6x increase in live iM-iPSCs (p<0.0001) and 12x in MG-iPSCs (p<0.0001) from first to last count. The average percentage of live cells for each day assessed was included in Fig 3C.

MTS assays were also performed and used as indicators of cell viability throughout 4 days in culture. This colorimetric assay is based on the reduction of a tetrazolium compound into a soluble colored formazan by metabolically active cells, which is proportional to the number of viable cells and the absorbance values obtained [18,19]. As represented in Fig 3D, MG-iPSCs showed significantly higher absorbance readings on days 2, 3, and 4. The most prominent difference was observed on day 2, with a 55% difference between groups (p=0.0003), which gradually decreased to 36% on day 3 (p=0.0045) and 29.4% on day 4 (p=0.0125). In contrast to the observed for the number of viable cells within ECMs, there was a significant increase in formazan conversion between days 1 and 2 (p=0.0285 for iM and p<0.0001 for MG) but no difference between the subsequent days.

### iM-iPSCs maintained pluripotency and genetic stability through 15 passages in culture

iMR90-4 iPSCs were thawed at P34 and grown up to P50 using either iM or MG. In all odd passages within this range cells were fixed for immunostainings and protein samples were collected. Fixed coverslips were used for phenotypic confirmation throughout 15 passages in culture. Expression of pluripotency related markers SSEA-4 and OCT-3/4 was successfully identified across passages for both iM- and MG-iPSCs. Representative images of cultures in P35 and P49, the first and last passages included in the study, are shown in Fig 4A and the complete panel containing all passages evaluated can be found in S1 Fig. Protein samples were used for the quantification of the phosphorylated form of histone H2AX (S139 γH2AX) through western blots. γH2AX is formed as a response to DNA double strand breaks (DSB) and is commonly used as a marker of DNA damage [20]. To establish a positive control, samples collected from iM-iPSCs in P48 irradiated at 2.14 Gy/min totalizing 8 Gy were included in each experiment and the averaged signal intensity of 12163.4 ± 4862.35 is represented by the red dotted line in Fig 4C. Šídák’s multiple comparisons test showed no statistical difference in γH2AX content in samples collected from iM- and MG-iPSCs across all passages, which was also shown to be equivalent across passages within each ECM through Tukey’s post-hoc test (Fig 4C). In fact, analysis of variance showed that ECM and passages accounted for only 5% (p=0.4057) and 14.5% (p=0.3285) of total variation, respectively. The lowest and highest relative signal intensities were observed at P47 (3629 ± 1031) and P43 (8300 ± 1031) for iM-iPSCs, and P39 (5835 ± 2717) and P45 (10904 ± 3834) for MG-iPSCs. No correlation between passage and γH2AX content was found for cells grown using either iM (p=0.2052, R^2^=0.2517) or MG (p=0.2855, R^2^=0.1864). The cumulative γH2AX content was calculated as the average relative signal intensity for all samples in each ECM and shown to be statistically equivalent (p=0.1251) between cells grown using iM (6356 ± 475.9) or MG (7812 ± 801.2), although variance was significant larger in the MG group (p=0.0155).

**Fig 4.**
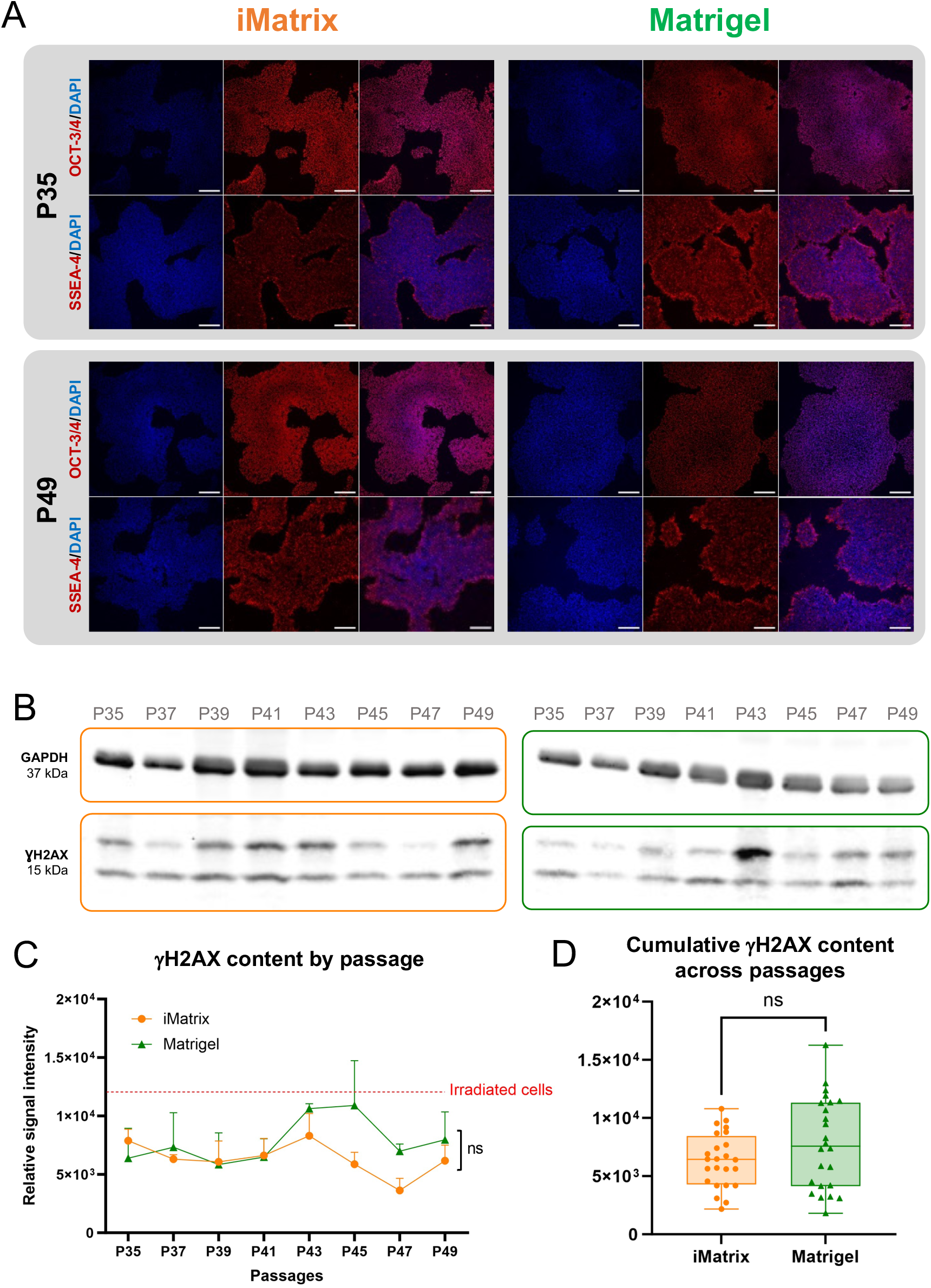
Pluripotency stability and DNA damage accumulation. **(A)** Confocal microscopy images of iM- and MG-iPSCs in passages 35 and 49 following immunolabeling of OCT-3/4 or SSEA-4 (red fluorescence) and nuclear staining (blue fluorescence). Scale bars: 200 µM. **(B)** Representative images of western blot membranes used for densitometric analysis to quantify γH2AX and GAPDH on protein samples obtained from 3 cultures of iM- and MG-iPSCs from passage 35 to 49. **(C)** Relative signal intensity over passages is represented as mean ± SEM. The red line represents mean signal intensity from irradiated iM-iPSCs. Statistical significance was assessed through two-way repeated measures ANOVA followed by Šídák’s and Tukey’s multiple comparison tests to compare iM- and MG-iPSCs in each passage and different passages within the same ECM group, respectively. n=3/passage/ECM. **(D)** Cumulative γH2AX content is represented as a Box & Whisker plot where the lines range from minimum to maximum values and the boxes depict the lower to upper quartile. Each point represents an individual measurement per passage, totalizing n=24/ECM. Statistical significance was evaluated through an unpaired t test. ns: not significant. DAPI: 4′,6-diamidino-2-phenylindole; GAPDH: glyceraldehyde 3-phosphate dehydrogenase; OCT-3/4: octamer binding transcription factor-3/4; P: passage; SSEA-4: stage-specific embryonic antigen-4; γH2AX: phosphorylated H2A histone family member X.

### iM supported directed differentiation of iPSCs into neurovascular cell types

iM- and MG-iPSCs were used to generate iBMECs and iPericytes following the equivalent ECM coating strategy according to the requirements of each differentiation protocol. Both target cell types were successfully generated using iM and MG, as shown in Figs 5 and 6. Images obtained after ZO-1 immunolabeling (Fig 5B) demonstrated that iM- and MG-iBMECs were capable of forming confluent monolayers and properly develop tight junctions. For further assessment of the development of barrier phenotypes by the iBMECs generated, TEER was measured over 5 days in culture. The obtained results revealed that resistance fluctuations followed a similar pattern for both cultures and no differences in average TEER was observed from day 1 to 4 after subculturing into transwell inserts, while statistical significance was found after 5 days in culture (p=0.0142, iM=54.88 ± 3.58 and MG=85.68 ± 7.03 Ω*cm^2^).

**Fig 5.**
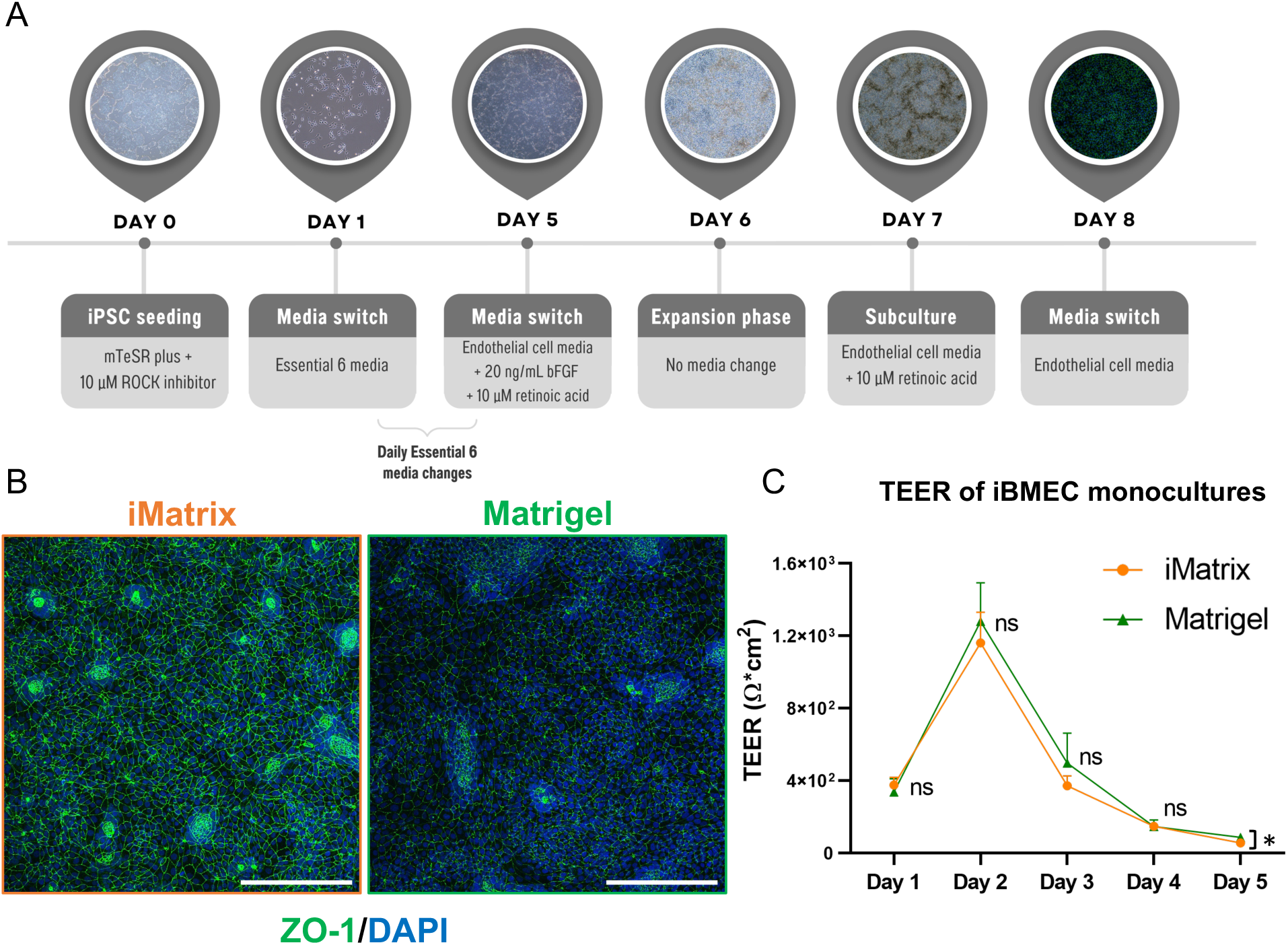
Generation of human iPSC-derived brain microvascular endothelial cells. **(A)** Schematic representation of the differentiation timeline. **(B)** Confocal microscopy images of iBMECs following immunolabeling of ZO-1 (green fluorescence) and nuclear staining (blue fluorescence). Scale bars: 200 µM. **(C)** Normalized TEER measurements over 5 days, represented as mean ± SEM, for demonstration of successful iBMEC differentiation through the quantification of barrier resistance. Statistical significance was assessed using a mixed effects model for repeated measures followed by Tukey’s multiple comparisons test. N=4-6 per time point measured in 3 independent experiments. *p=0.0142, ns: not significant. DAPI: 4′,6-diamidino-2-phenylindole; iBMEC: iPSC-derived brain microvascular endothelial cells; TEER: transendothelial electrical resistance; ZO-1: zonula occludens-1.

Normalized resistance peaked after 48h in culture for both groups, reaching 1161 ± 168.3 03 Ω*cm^2^ for iM-iBMECs and 1279 ± 213.8 03 Ω*cm^2^ for MG-iBMECs (p=0.6738). Peak TEER represented a 102.4% increase from 374.8 ± 43.49 Ω*cm^2^ on day 1 to 1161 ± 168.3 Ω*cm^2^ on day 2 for iM-iBMECs (p=0.0119), and a 116.5% increase from 337.3 ± 72.41 Ω*cm^2^ to 1279 ± 213.8 Ω*cm^2^ for their MG counterparts (p=0.0282). A similar trend was observed in the following drop in resistance from day 2 to 3. While iM-BMECs showed a 103% decrease to 371.5 ± 54.92 Ω*cm^2^ (p=0.0078), an 88.2% decrease to 496.3 ± 165.3 Ω*cm^2^ (p=0.0064) was observed for MG-iBMECs.

It has been suggested that although most peripheral pericytes have a mesodermal origin, neural crest stem cells can be the developmental origin of the pericytes found in the central nervous system, which can potentially entail important functional distinctions [21]. Therefore, a methodological approach in which iPSCs are first differentiated into neural crest stem cells (iNCSC) before being induced into iPericytes was adopted.

As shown by the expression of cell specific markers (Fig 6), we were able to successfully differentiate iMR90-4 iPSCs into iNCSC and the latter into iPericytes using both matrices. Projected Z-stacks of the immunoidentification of NGFR in non-sorted iNCSC showed extensive expression of this marker across groups (Fig 6B), and on average the cell sorting process yielded 174,792 ± 14,941 and 168,958 ± 23,275 NGFR^+^ cells per cm^2^ collected for iNCSC generated using iM and MG, respectively. An unpaired t test showed that the mean yield (p=0.8433) and variances (p=0.5836) were statistically equivalent (n=3). Furthermore, acquisition of expression of NG-2 and PDGFR-β by NGFR^+^ cells after 9 days of culture in E6 + 10% FBS on non-coated surfaces demonstrates that fully differentiated iPericytes can be obtained through the proposed method using either coating strategy (Fig 6C).

**Fig 6.**
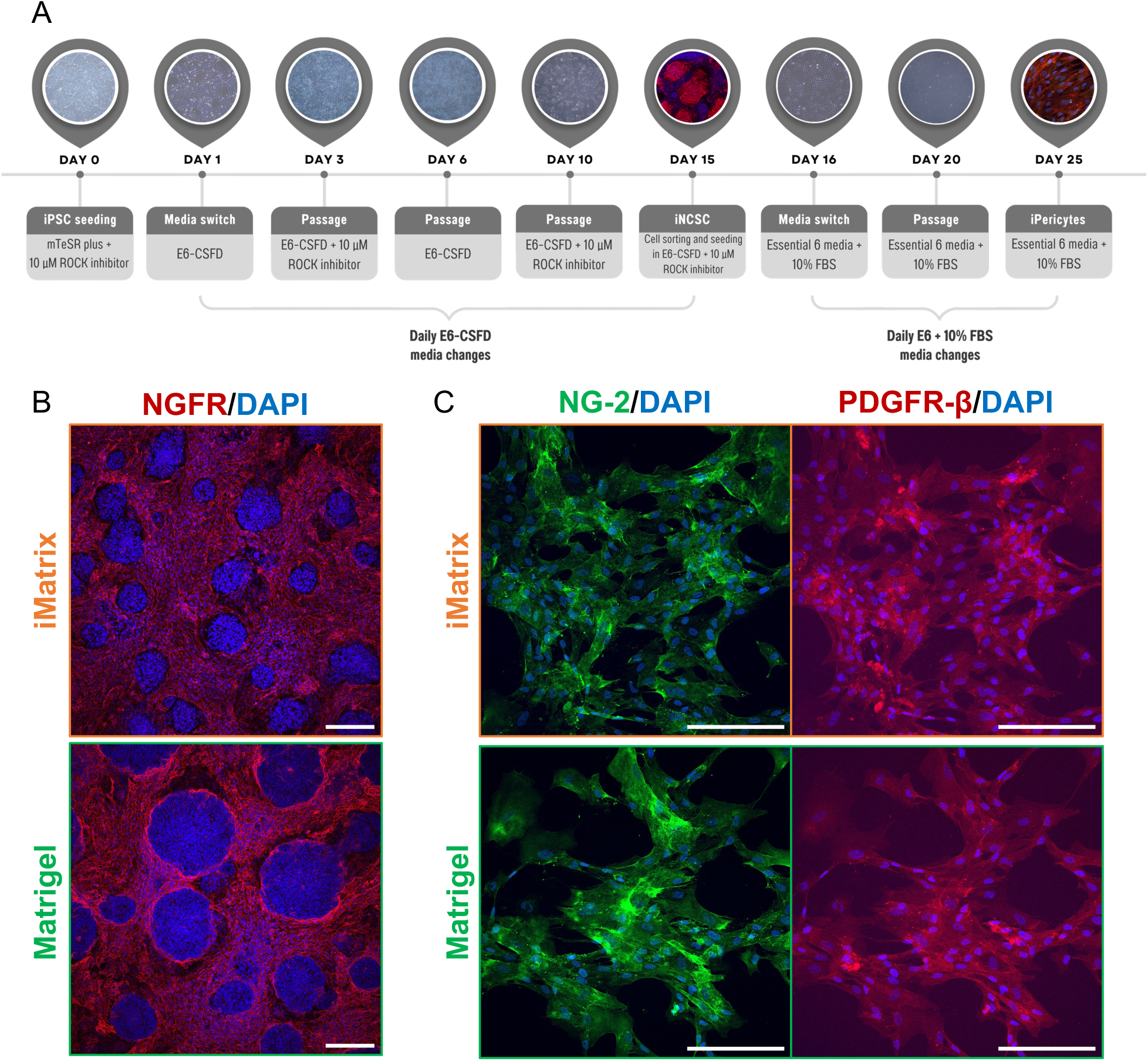
Generation of human iPSC-derived brain pericytes. **(A)** Schematic representation of the differentiation timeline. **(B)** Confocal microscopy images of iNCSCs following immunolabeling of NGFR (red fluorescence) and nuclear staining (blue fluorescence). **(C)** Confocal microscopy images of fully differentiated iPericytes following immunolabeling of PDGFR-β (green fluorescence), NG-2 (red fluorescence), and nuclear staining (blue fluorescence). Scale bars: 200 µM. DAPI: 4′,6-diamidino-2-phenylindole; E6-CSFD: Essential 6 media supplemented with CHIR99021, SB431542, FGF-2, dorsomorphin, and heparin; FBS: fetal bovine serum; NG-2: neural/glial antigen 2; NGFR: nerve growth factor receptor; PDGFR-β: platelet-derived growth factor receptor β.

One important drawback of the application of iPericytes for BBB modeling in this context, is the length of the differentiation process. While iBMEC can be generated and seeded for experimental analyses in 8 days, the differentiation of iPSCs into iPericytes is a 25-days long process, as shown in the diagrams on Figs 5A and 6A. This factor hampers experimental planning and requires that several simultaneous differentiation processes are performed to reduce the interval between successive experimental rounds. Aiming to overcome this limitation, we investigated the possibility to cryopreserve the cells in two different stages of differentiation. Initially, iNCSC were frozen in serum free cell freezing media (Sigma) immediately after sorting to take advantage of the addition of ROCK inhibitor to the cell culture media for 24 hours in this step of the process. However, cells did not reattach when thawed in E6+20% FBS onto non-coated cell culture plates.

The second approach tested was the cryopreservation of fully differentiated iPericytes after day 25 in a freezing media composed of 50% E6, 40% FBS, and 10% dimethyl sulfoxide (DMSO; Sigma) in the presence and absence of ROCK inhibitor. iPericytes were frozen at a density of 1x10^6^ cells per cryovial and for thawing 1 vial was seeded onto a non-coated T-25 cell culture flask (Corning) in E6+20% FBS, and the serum concentration was decreased to 10% after 24 hours. In this case, cell recovery was successful after maintenance in liquid nitrogen, and more efficient for cells frozen with ROCK inhibitor, which showed improved cell adhesion and survival in the first 24 hours following seeding (S2 Fig B). Phenotyping of thawed cells demonstrated that the expression of cell specific markers was retained after freezing (S2 Fig A). The successful cryopreservation of iPericytes described here represents an important tool to enhance experimental efficiency for human iPSC-based isogenic multicellular BBB modeling.

### iM serves as a suitable substrate for generation of iPSC-derived cells for BBB modeling

iBMECs and iPericytes were paired by ECM and co-cultured on a transwell system, as demonstrated in Fig 7A, for a comparative assessment of their performance as a BBB model following the experimental timeline shown in Fig 7B. Co-cultures were maintained for 5 days and throughout this period TEER, permeability, and expression of tight junction proteins were measured as indicators of development of barrier properties. When it comes to TEER, no significant differences were observed between iM and MG co-cultures from day 1 to 5, as shown in Fig 7C. Peak resistance was achieved 48h after seeding for co-cultures in both groups, reaching 1410 ± 91.67 Ω*cm^2^ for iM and 1265 ± 121.7 Ω*cm^2^ for MG, representing a 137% (p=0.0131) and 118.5% (p=0.5656) increase from day 1, respectively. The decrease in TEER observed between days 2 and 3 was not significant for either group (p=0.0666 for iM and p=0.1704 for MG). After 5 days in culture, resistance significantly dropped compared to peak values for both iM (p=0.0097) and MG (p=0.03) co-cultures.

**Fig 7.**
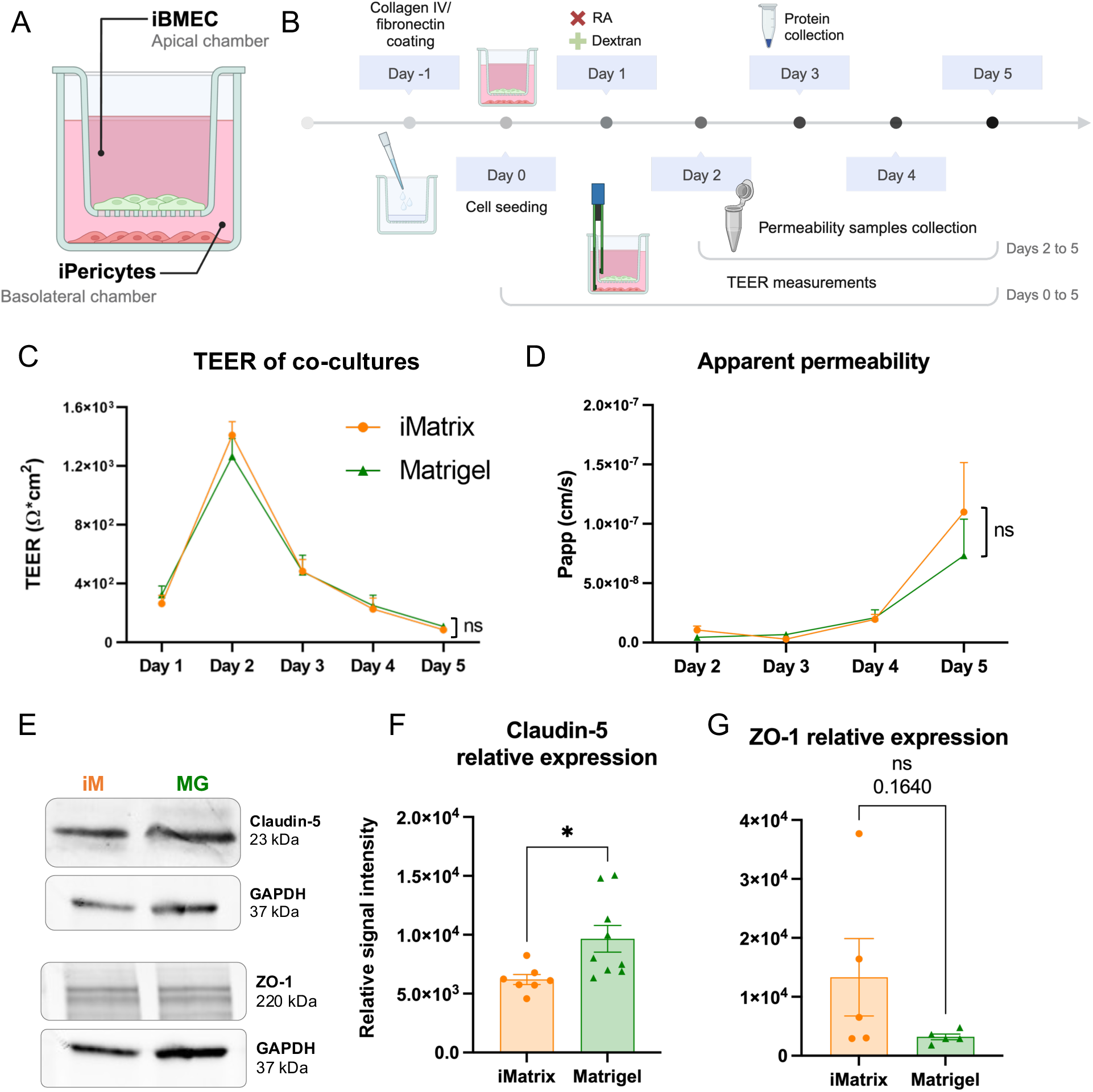
Co-culture of isogenic iPSC-derived brain microvascular endothelial cells and brain pericytes. **(A)** Schematic representation of co-cultures on transwells, showing iBMECs on the apical chamber and iPericytes on the basolateral chamber. **(B)** Representative diagram of the experimental timeline adopted for co-culture studies. **(C)** Normalized TEER measurements over 5 days, represented as mean ± SEM. Statistical significance of averaged replicates per time point was evaluated through two-way ANOVA followed by Tukey’s multiple comparisons test. N=6-9 per time point measured in 3 independent experiments. **(D)** Apparent permeability of iBMECs monolayers to 3 kDa dextran over 4 days in culture represented as mean ± SEM. Statistical significance was assessed using a mixed effects model for repeated measures followed by Tukey’s multiple comparisons test. N=4-6 per time point measured in 3 independent experiments. **(E)** Expression of tight junction proteins was evaluated using western blots. Relative signal intensity of **(F)** claudin-5 and **(G)** ZO-1 was determined through densitometric analysis using GAPDH expression for normalization and plotted as mean ± SEM. Significance of mean differences was assessed through unpaired t tests with a total n=7-9 for claudin-5 and n=5 for ZO-1 corresponding to samples collected in 3 independent experiments. *p=0.022, ns: not significant. GAPDH: glyceraldehyde 3-phosphate dehydrogenase; iBMEC: iPSC-derived brain microvascular endothelial cells; iM: iMatrix; MG: Matrigel; Papp: apparent permeability; RA: retinoic acid; TEER: transendothelial electrical resistance; ZO-1: zonula occludens 1. A & B were created using BioRender.

Apparent permeability was also found to be statistically equivalent between iM- and MG-derived co-cultures in all time points included in this study (Fig 7D). In a closer look, a significant increase in permeability was observed between days 2 and 5 for both experimental groups. iM-derived co-cultures showed a 10.5x increase in this period (p=0.0021), while for their MG counterparts a 16.7x difference was observed (p=0.0439). Co-cultures generated using iM-derived cells also showed significantly higher permeability on day 5 compared to days 3 (p=0.0081) and 4 (p=0.0054), while the same was not observed for MG-derived cells.

Finally, expression of tight junction proteins claudin-5 and ZO-1 was analyzed through western blots (Fig 7E-G). On day 3, MG-iBMECs (9660 ± 1129) in co-culture showed significantly higher expression of claudin-5 (p=0.0220) compared to iM-iBMECs (6204 ± 424.2). Variance was also found to be higher for MG-iBMECs (p=0.0148) (Fig 7F). ZO-1 expression, on the other hand, was equivalent between both groups (p=0.1640), although variance in this case was significantly higher for iM-iBMECs (p=0.0002) (Fig 7G).

## Discussion

This study investigated the use of iM, a commercially available recombinant fragment of human laminin-511, as a xenofree and fully defined alternative to MG. While earlier studies have demonstrated that the fragment E8 of laminin-511 (LN511-E8) supports maintenance and differentiation of pluripotent stem cells in vitro [2,9,22,23], its use to reproduce methods validated using MG have not been explicitly addressed. In this study, we have demonstrated that iM-iPSCs behave similarly to MG-iPSCs and can be differentiated into neurovascular cell types using protocols established using MG.

A key requirement of iPSC maintenance is that these cells require passaging in aggregates to avoid myosin-actin contraction related cell death without the need to incorporate ROCK inhibitor in routine expansion protocols. Ideally, aggregates should be consistent in size while large enough to reattach as colonies and ensure cell survival and small enough to facilitate access by growth factors and other small molecules [24,25]. Through colony attachment analysis, we found that 24h after seeding iM-iPSCs are significantly less variable in size than MG-iPSCs and, although not significantly, smaller considering the surface area covered. Consistently, more colonies were formed upon attachment to iM, suggesting that iPSCs grown on iM reattach in more colonies that are smaller in size compared to MG-cultured cells, which formed more and larger colonies. These observations can be associated with the consistency of cell dissociation for expansion and the size and number of aggregates generated as cells anchored to both matrices are released from the plate and neighboring cells. Therefore, iM might be more compatible with aggregate passaging [4,24]. A previous study has also demonstrated that LN511-E8 support stronger adhesion of singularized iPSCs than MG and even intact LN511, while making the addition of ROCK inhibitor optional following complete dissociation and allowing for uniform cell seeding in defined densities [2].

When it comes to cell behavior in culture, we have demonstrated that the number of iM-iPSCs and MG-iPSCs increased similarly in the typical interval between passages [3], showing stable and consistent activation of DNA damage responses and retaining expression of pluripotency markers across 15 passages. Interestingly, while the number of live and dead cells after trypan-blue staining was statistically equivalent for iM and MG in all time points, MTS readings were significantly higher for MG-iPSCs from day 2 to 4 after seeding. Note that, although not significantly, live cell counts were higher for iM-iPSCs every day, with the smallest difference coinciding with the higher difference in MTS readings on day 2. Interestingly, these two methods rely on different cellular responses. While trypan blue allows for the identification of cells with compromised membranes, the MTS assay detects differences in mitochondrial metabolism as the reaction quantified is believed to be performed mainly by NAD(P)H-dependent oxidoreductases [26,27].

In a study comparing four methods for determining viability of fibroblasts after irradiation with increasing doses of blue light, statistical significance was first detected with a tetrazolium-based assay (i.e. MTT) at 55 J/cm^2^ and subsequently with MTT and trypan blue at 110 J/cm^2^. The reported data demonstrates that mitochondrial metabolism is affected before membrane integrity is compromised, demonstrating that these methods differ in sensitivity [27]. In this case, the difference we observed could be a result of reduced metabolic activity in iPSCs grown on iM that did not reflect in membrane damage. On the other hand, there is evidence that metabolic dysregulation may increase the flow of electrons to artificial dyes through compensatory reactions, which can be mistaken by increased NAD(P)H available for cell proliferation [26]. Although the effects of MG on iPSC metabolism have not yet been directly addressed, it has been shown that its incorporation into the media for culture of breast cancer spheroids under hypoxia leads to increased Nrf2 expression and nitric oxide levels, as well as decreased levels of glutathione compared to cells kept in absence of MG, therefore attesting for a potential role of its bioactive components in modulating cellular metabolism [28]. It is also important to take into account that MG was prepared for surface coating in DMEM/F12 containing glucose, where its presence in culture media has been shown to increase tetrazolium reduction in HEK293 and HeLa cells [29]. Overall, the obtained data suggests that iM supports iPSCs growth and survival similarly to MG. These results also emphasize the need for future studies providing a closer look at possible effects of MG in modulating mitochondrial metabolism given its major role in pluripotency maintenance [30] and the importance of combining different viability assays for a better understanding of cellular responses [27].

According to evidence showing that the full form of laminin-511 [8] and LN511-E8 [2,9] support continuous subculture of pluripotent stem cells with normal karyotypes for up to 30 passages, our analysis of γH2AX content in cells grown on both matrices suggests that iM works as well as MG when it comes to maintaining genetic stability across passages. Considered the most sensitive marker for detection of DNA damage, γH2AX is the phosphorylated form of histone H2AX, which is rapidly formed in response to DNA double-strand breaks [31]. It is important to note that iPSCs show a higher basal level of γH2AX compared to somatic cell types, which has been suggested to be associated with increased proliferation rates and shown to decrease in differentiated progeny [32].

In addition to genetic stability, previous studies have demonstrated that LN511-E8 supports long-term maintenance of undifferentiated iPSCs with steady expression of several pluripotency markers [2,9,22], including SSEA-4 and OCT-3/4, which were also observed in cultures grown on both iM and MG in every odd passage between 35 and 49 in this study. Tied to proper maintenance of pluripotency is the ability of cells to respond properly to stimuli to differentiate into somatic cell types – a critical step for most iPSCs applications. Several differentiation protocols are performed using MG-coated surfaces, thus an efficient ECM alternative would ideally be completely interchangeable, supporting targeted differentiation as well as undifferentiated proliferation. It has been shown that iPSCs cultured on LN511-E8 can generate embryoid bodies and teratomas expressing gene markers for all three germ layers [2,9,22] and undergo directed differentiation into dopaminergic neurons, blood cells, and insulin-producing cells with LN511-E8 being incorporated in steps that required ECM coating [22]. To complement what is known about the compatibility of LN511-E8 with differentiation protocols, we have demonstrated that methods previously validated using MG for the generation of iBMECs, iNCSCs, and iPericytes [14,15] can be replicated using iM.

iPSC-derived cells are a powerful tool for disease modeling, impacting immensely the physiological relevance of benchtop models based on their applicability for personalized medicine and development of increasingly complex isogenic multicellular systems [33]. Therefore, following successful generation of neurovascular cell types using iM and MG, isogenic iBMECs and iPericytes generated using the same ECM were combined and co-cultured [15] for 5 days in a transwell system for a comparative assessment of the development of barrier phenotypes. Characterization of co-cultures of iPSC derived BMECs and pericytes and even BMEC monocultures have shown that cellular behavior can differ substantially between studies, for example reaching higher [34] and lower [15] TEER values than reported here, which is possibly a consequence of significant variations in differentiation protocols and iPSC lines used. Nonetheless, our results demonstrate that iBMECs generated using both strategies can form proper tight junctions [34,35] and peak in resistance above 1000 Ω*cm^2^ after 48h in co-culture with iPericytes. As expected, the highest apparent permeability coincided with the lowest TEER values on day 5 in both conditions, suggesting the loss of barrier integrity over time. Other studies have shown that iPSC-derived BBB models can reach TEER values up to 4000 Ω*cm^2^ [14,34]. Differences in TEER numbers between these previous studies and our current work are possibly due to differences in specific culture reagents used. However, as cells from both treatment groups were manipulated simultaneously and exposed to the exact same conditions in each experiment, these differences in TEER numbers from previous studies are not detrimental to a comparative analysis.

Finally, complementing the assessment of BBB-like phenotypes in endothelial cells, we have demonstrated using a semiquantitative method that the expression of claudin-5 is upregulated in MG-iBMECs compared to iM-iBMECs, while ZO-1 expression was statistically equivalent between both groups despite a trend towards higher expression by iM-iBMECs. Proper formation of tight junctions is essential for the establishment of inter-endothelial connections, which plays a major role in BBB tightness and healthy function, controlling paracellular transport and barrier permeability [36]. Several studies have previously demonstrated that iBMECs generated using MG or vitronectin express cell junction proteins, such as VE-cadherin, occludin, claudin-5, and ZO-1 [14,15,34,37]. Claudin-5, the most enriched tight junction protein in the neurovasculature, is an integral membrane protein that tightly connects adjacent cells, and its intracellular domains are anchored to the actin cytoskeleton through scaffolding proteins such as ZO-1/2/3 [36]. Its role in controlling barrier permeability is so important, that its selective modulation is being studied as a pharmacological target to increase drug delivery to the brain and as a strategy to stabilize cellular junctions to treat disorders associated with BBB leakage [38]. Similarly, it has been demonstrated that cells lacking ZO proteins show disrupted tight junction formation, with claudins failing to polymerize [39], hence the importance of demonstrating that iM- and MG-iBMECs express both proteins. Taken together, the presented data suggests that although MG-iBMECs show higher claudin-5 expression, the observed difference did not result in functional changes in barrier tightness under evaluated conditions, as confirmed by statistically equivalent TEER and apparent permeability through 5 days in co-culture with iPericytes. In this case, if also taking into account the presence of known and unknown bioactive components in MG, the possibility of its effect on differences in gene and protein expression cannot be overlooked [4] and should be addressed in future studies.

Overall, our findings indicate that iM is a viable fully defined and xenofree alternative to MG for iPSC maintenance and differentiation into relevant cell types for neurovascular modeling, yielding similar results in the establishment of a simplified isogenic iPSC-derived BBB in vitro system. Ultimately, evidence presented here will contribute to increasing flexibility in the development of protocols for culture and targeted differentiation of pluripotent stem cells, while supporting the enhanced experimental robustness and reproducibility that comes with the adoption of fully defined and batch-to-batch consistent ECM products for cell culture.

## Acknowledgments

The authors would like to thank the contributions of Dr. Young Hye Song and In Ha Baek (University of Arkansas), for providing training and access to their confocal microscope for acquisition of all ICC images included in this publication; and Dr. Rajaram and Dr. Whitney Ann Lowe (University of Arkansas) for enabling and assisting with the irradiation of iPSC cultures.

## References

1. Thomson JA, Itskovitz-Eldor J, Shapiro SS, Waknitz MA, Swiergiel JJ, Marshall VS, et al. Embryonic Stem Cell Lines Derived from Human Blastocysts. Science [Internet]. 1998 Nov 6 [cited 2024 May 13];282(5391):1145–7. Available from: https://www.science.org/doi/10.1126/science.282.5391.1145

2. Miyazaki T, Futaki S, Suemori H, Taniguchi Y, Yamada M, Kawasaki M, et al. Laminin E8 fragments support efficient adhesion and expansion of dissociated human pluripotent stem cells. Nat Commun [Internet]. 2012 Dec 4 [cited 2024 Apr 28];3(1):1236. Available from: https://www.nature.com/articles/ncomms2231

3. Lyra-Leite DM, Gutiérrez-Gutiérrez Ó, Wang M, Zhou Y, Cyganek L, Burridge PW. A review of protocols for human iPSC culture, cardiac differentiation, subtype-specification, maturation, and direct reprogramming. STAR Protocols [Internet]. 2022 Sep [cited 2024 May 5];3(3):101560. Available from: https://linkinghub.elsevier.com/retrieve/pii/S2666166722004403

4. Aisenbrey EA, Murphy WL. Synthetic alternatives to Matrigel. Nat Rev Mater [Internet]. 2020 Jul [cited 2021 Nov 15];5(7):539–51. Available from: http://www.nature.com/articles/s41578-020-0199-8

5. Vukicevic S, Kleinman HK, Luyten FP, Roberts AB, Roche NS, Reddi AH. Identification of multiple active growth factors in basement membrane matrigel suggests caution in interpretation of cellular activity related to extracellular matrix components. Experimental Cell Research [Internet]. 1992 Sep 1 [cited 2024 May 3];202(1):1–8. Available from: https://www.sciencedirect.com/science/article/pii/001448279290397Q

6. Kiyozumi D, Nakano I, Sato-Nishiuchi R, Tanaka S, Sekiguchi K. Laminin is the ECM niche for trophoblast stem cells. Life Science Alliance [Internet]. 2020 Feb 1 [cited 2024 May 13];3(2). Available from: https://www.life-science-alliance.org/content/3/2/e201900515

7. Miyazaki T, Futaki S, Hasegawa K, Kawasaki M, Sanzen N, Hayashi M, et al. Recombinant human laminin isoforms can support the undifferentiated growth of human embryonic stem cells. Biochemical and Biophysical Research Communications [Internet]. 2008 Oct [cited 2024 May 13];375(1):27–32. Available from: https://linkinghub.elsevier.com/retrieve/pii/S0006291X08014319

8. Rodin S, Domogatskaya A, Ström S, Hansson EM, Chien KR, Inzunza J, et al. Long-term self-renewal of human pluripotent stem cells on human recombinant laminin-511. Nat Biotechnol [Internet]. 2010 Jun [cited 2024 Apr 28];28(6):611–5. Available from: https://www.nature.com/articles/nbt.1620

9. Miyazaki T, Isobe T, Nakatsuji N, Suemori H. Efficient Adhesion Culture of Human Pluripotent Stem Cells Using Laminin Fragments in an Uncoated Manner. Sci Rep [Internet]. 2017 Mar [cited 2021 Oct 14];7(1):41165. Available from: http://www.nature.com/articles/srep41165

10. Kuo HH, Gao X, DeKeyser JM, Fetterman KA, Pinheiro EA, Weddle CJ, et al. Negligible-Cost and Weekend-Free Chemically Defined Human iPSC Culture. Stem Cell Reports [Internet]. 2020 Feb [cited 2023 Dec 9];14(2):256–70. Available from: https://linkinghub.elsevier.com/retrieve/pii/S2213671119304461

11. Shi G, Jin Y. Role of Oct4 in maintaining and regaining stem cell pluripotency. Stem Cell Research & Therapy [Internet]. 2010 Dec 14 [cited 2024 Feb 2];1(5):39. Available from: 10.1186/scrt39

12. Virant-Klun I, Kenda-Suster N, Smrkolj S. Small putative NANOG, SOX2, and SSEA-4-positive stem cells resembling very small embryonic-like stem cells in sections of ovarian tissue in patients with ovarian cancer. J Ovarian Res [Internet]. 2016 Mar 3 [cited 2024 Feb 2];9:12. Available from: https://www.ncbi.nlm.nih.gov/pmc/articles/PMC4778328/

13. Ghasemi M, Turnbull T, Sebastian S, Kempson I. The MTT Assay: Utility, Limitations, Pitfalls, and Interpretation in Bulk and Single-Cell Analysis. IJMS [Internet]. 2021 Nov 26 [cited 2023 Dec 9];22(23):12827. Available from: https://www.mdpi.com/1422-0067/22/23/12827

14. Neal EH, Marinelli NA, Shi Y, McClatchey PM, Balotin KM, Gullett DR, et al. A Simplified, Fully Defined Differentiation Scheme for Producing Blood-Brain Barrier Endothelial Cells from Human iPSCs. Stem Cell Reports [Internet]. 2019 Jun [cited 2020 Aug 29];12(6):1380–8. Available from: https://linkinghub.elsevier.com/retrieve/pii/S2213671119301754

15. Stebbins MJ, Gastfriend BD, Canfield SG, Lee MS, Richards D, Faubion MG, et al. Human pluripotent stem cell–derived brain pericyte–like cells induce blood-brain barrier properties. Sci Adv [Internet]. 2019 Mar 20 [cited 2021 Oct 14];5(3):eaau7375. Available from: https://www.science.org/doi/10.1126/sciadv.aau7375

16. McCallinhart PE, Biwer LA, Clark OE, Isakson BE, Lilly B, Trask AJ. Myoendothelial Junctions of Mature Coronary Vessels Express Notch Signaling Proteins. Front Physiol [Internet]. 2020 Feb 4 [cited 2023 Dec 10];11:29. Available from: https://www.frontiersin.org/article/10.3389/fphys.2020.00029/full

17. Piña R, Santos-Díaz AI, Orta-Salazar E, Aguilar-Vazquez AR, Mantellero CA, Acosta-Galeana I, et al. Ten Approaches That Improve Immunostaining: A Review of the Latest Advances for the Optimization of Immunofluorescence. IJMS [Internet]. 2022 Jan 26 [cited 2023 Dec 10];23(3):1426. Available from: https://www.mdpi.com/1422-0067/23/3/1426

18. Berridge MV, Herst PM, Tan AS. Tetrazolium dyes as tools in cell biology: New insights into their cellular reduction. In Biotechnology Annual Review [Internet]. Elsevier; 2005 [cited 2024 Jan 21]. p. 127–52. Available from: https://linkinghub.elsevier.com/retrieve/pii/S1387265605110047

19. Stepanenko AA, Dmitrenko VV. Pitfalls of the MTT assay: Direct and off-target effects of inhibitors can result in over/underestimation of cell viability. Gene [Internet]. 2015 Dec [cited 2024 Jan 21];574(2):193–203. Available from: https://linkinghub.elsevier.com/retrieve/pii/S037811191500952X

20. Mariotti LG, Pirovano G, Savage KI, Ghita M, Ottolenghi A, Prise KM, et al. Use of the γ-H2AX Assay to Investigate DNA Repair Dynamics Following Multiple Radiation Exposures. Robson CN, editor. PLoS ONE [Internet]. 2013 Nov 29 [cited 2024 Feb 2];8(11):e79541. Available from: https://dx.plos.org/10.1371/journal.pone.0079541

21. Jamieson JJ, Searson PC, Gerecht S. Engineering the human blood-brain barrier in vitro. J Biol Eng [Internet]. 2017 Dec [cited 2022 Jul 31];11(1):37. Available from: https://jbioleng.biomedcentral.com/articles/10.1186/s13036-017-0076-1

22. Nakagawa M, Taniguchi Y, Senda S, Takizawa N, Ichisaka T, Asano K, et al. A novel efficient feeder-free culture system for the derivation of human induced pluripotent stem cells. Sci Rep [Internet]. 2014 Jan 8 [cited 2024 Apr 28];4(1):3594. Available from: https://www.nature.com/articles/srep03594

23. Nakashima Y, Tsukahara M. Laminin-511 Activates the Human Induced Pluripotent Stem Cell Survival via α6β1 Integrin-Fyn-RhoA-ROCK Signaling. Stem Cells Dev [Internet]. 2022 Nov 1 [cited 2024 Apr 30];31(21–22):706–19. Available from: https://www.ncbi.nlm.nih.gov/pmc/articles/PMC9700348/

24. Beers J, Gulbranson DR, George N, Siniscalchi LI, Jones J, Thomson JA, et al. Passaging and colony expansion of human pluripotent stem cells by enzyme-free dissociation in chemically defined culture conditions. Nat Protoc [Internet]. 2012 Nov [cited 2024 Apr 26];7(11):2029–40. Available from: https://www.nature.com/articles/nprot.2012.130

25. Chen G, Hou Z, Gulbranson D, Thomson JA. Actin-myosin contractility is responsible for the reduced viability of dissociated human embryonic stem cells. Cell Stem Cell [Internet]. 2010 Aug 6 [cited 2024 Apr 26];7(2):240–8. Available from: https://www.ncbi.nlm.nih.gov/pmc/articles/PMC2916864/

26. Aleshin VA, Artiukhov AV, Oppermann H, Kazantsev AV, Lukashev NV, Bunik VI. Mitochondrial Impairment May Increase Cellular NAD(P)H: Resazurin Oxidoreductase Activity, Perturbing the NAD(P)H-Based Viability Assays. Cells [Internet]. 2015 Aug 21 [cited 2024 Apr 29];4(3):427–51. Available from: https://www.ncbi.nlm.nih.gov/pmc/articles/PMC4588044/

27. Masson-Meyers DS, Bumah VV, Enwemeka CS. A comparison of four methods for determining viability in human dermal fibroblasts irradiated with blue light. Journal of Pharmacological and Toxicological Methods [Internet]. 2016 May [cited 2024 Apr 29];79:15–22. Available from: https://linkinghub.elsevier.com/retrieve/pii/S1056871916300016

28. Badea MA, Balas M, Hermenean A, Ciceu A, Herman H, Ionita D, et al. Influence of Matrigel on Single- and Multiple-Spheroid Cultures in Breast Cancer Research. SLAS Discovery [Internet]. 2019 Jun 1 [cited 2024 Apr 29];24(5):563–78. Available from: https://www.sciencedirect.com/science/article/pii/S247255522206782X

29. Xie L, Dai Z, Pang C, Lin D, Zheng M. Cellular glucose metabolism is essential for the reduction of cell-impermeable water-soluble tetrazolium (WST) dyes. Int J Biol Sci [Internet]. 2018 Sep 7 [cited 2024 May 12];14(11):1535–44. Available from: https://www.ncbi.nlm.nih.gov/pmc/articles/PMC6158726/

30. Xu X, Duan S, Yi F, Ocampo A, Liu GH, Izpisua Belmonte JC. Mitochondrial Regulation in Pluripotent Stem Cells. Cell Metabolism [Internet]. 2013 Sep [cited 2024 Apr 29];18(3):325–32. Available from: https://linkinghub.elsevier.com/retrieve/pii/S1550413113002490

31. Zhao H, Zhuang Y, Li R, Liu Y, Mei Z, He Z, et al. Effects of different doses of X-ray irradiation on cell apoptosis, cell cycle, DNA damage repair and glycolysis in HeLa cells. Oncol Lett [Internet]. 2019 Jan [cited 2024 May 12];17(1):42–54. Available from: https://www.ncbi.nlm.nih.gov/pmc/articles/PMC6313204/

32. Vallabhaneni H, Lynch PJ, Chen G, Park K, Liu Y, Goehe R, et al. High Basal Levels of γH2AX in Human Induced Pluripotent Stem Cells Are Linked to Replication-Associated DNA Damage and Repair. Stem Cells [Internet]. 2018 Oct [cited 2024 May 12];36(10):1501–13. Available from: https://www.ncbi.nlm.nih.gov/pmc/articles/PMC6662168/

33. Sharma A, Sances S, Workman MJ, Svendsen CN. Multi-lineage Human iPSC-Derived Platforms for Disease Modeling and Drug Discovery. Cell Stem Cell [Internet]. 2020 Mar [cited 2024 May 5];26(3):309–29. Available from: https://linkinghub.elsevier.com/retrieve/pii/S1934590920300643

34. Jamieson JJ, Linville RM, Ding YY, Gerecht S, Searson PC. Role of iPSC-derived pericytes on barrier function of iPSC-derived brain microvascular endothelial cells in 2D and 3D. Fluids Barriers CNS [Internet]. 2019 Dec [cited 2020 Sep 15];16(1):15. Available from: https://fluidsbarrierscns.biomedcentral.com/articles/10.1186/s12987-019-0136-7

35. Noorani B, Bhalerao A, Raut S, Nozohouri E, Bickel U, Cucullo L. A Quasi-Physiological Microfluidic Blood-Brain Barrier Model for Brain Permeability Studies. Pharmaceutics [Internet]. 2021 Sep 15 [cited 2024 May 12];13(9):1474. Available from: https://www.ncbi.nlm.nih.gov/pmc/articles/PMC8468926/

36. Greene C, Hanley N, Campbell M. Claudin-5: gatekeeper of neurological function. Fluids and Barriers of the CNS [Internet]. 2019 Jan 29 [cited 2024 May 13];16(1):3. Available from: 10.1186/s12987-019-0123-z

37. Gopinadhan A, Hughes JM, Conroy AL, John CC, Canfield SG, Datta D. A human pluripotent stem cell-derived in vitro model of the blood–brain barrier in cerebral malaria. Fluids Barriers CNS [Internet]. 2024 May 1 [cited 2024 May 13];21(1):38. Available from: https://fluidsbarrierscns.biomedcentral.com/articles/10.1186/s12987-024-00541-9

38. Hashimoto Y, Campbell M, Tachibana K, Okada Y, Kondoh M. Claudin-5: A Pharmacological Target to Modify the Permeability of the Blood–Brain Barrier. Biological & Pharmaceutical Bulletin [Internet]. 2021 Oct 1 [cited 2024 May 13];44(10):1380–90. Available from: https://www.jstage.jst.go.jp/article/bpb/44/10/44_b21-00408/_article

39. Liu W, Wang Z, Zhang L, Wei X, Li L. Tight Junction in Blood-Brain Barrier: An Overview of Structure, Regulation, and Regulator Substances. CNS Neuroscience & Therapeutics [Internet]. 2012 Aug [cited 2024 May 13];18(8):609–15. Available from: https://onlinelibrary.wiley.com/doi/10.1111/j.1755-5949.2012.00340.x

