## Supplemental Documents for "The future is fully defined: recombinant fragment E8 of laminin-511 is a viable xenofree alternative to Matrigel for hiPSC culture and differentiation into neurovascular cell types"

**Supplemental Table 1.** Reagents used for cell culture and assays.

| Product | Vendor | Catalog number |
|---|---|---|
| 3 kDa Cascade Blue-Dextran | Invitrogen | D7132 |
| B27 | Thermo Fisher Scientific | 17504044 |
| bFGF/FGF-2 | Preprotech | 100-18B |
| Bovine Serum Albumin | Millipore Sigma | A9418 |
| CHIR99021 | Sigma-Aldrich | SML1046 |
| Collagen IV from Human Placenta | Sigma-Aldrich | C5533 |
| DAPI (4',6-diamidino-2-phenylindole) | Thermo Fisher | D1306 |
| DMEM/F-12 | Thermo Fisher Scientific | 11330-032 |
| Dorsomorphin | Tocris | 3093 |
| Essential 6 Media | Thermo Fisher Scientific | A1516401 |
| Ethylenediaminetetraacetic acid (EDTA) | BioRad | 161-0729 |
| FcR Blocking Agent | Miltenyi Biotec | 130-059-901 |
| Fetal Bovine Serum | Thermo Fisher Scientific | 26140079 |
| Fibronectin from Bovine Plasma | Sigma-Aldrich | F1141 |
| Gelatin from cold water fish skin | Sigma-Aldrich | G7041 |
| Growth Factor Reduced Matrigel | Corning | 47743-718 |
| Heparin Sodium Salt from Porcine Intestinal Mucosa | Sigma-Aldrich | H3149 |
| Human Endothelial Serum Free Medium | Thermo Fisher Scientific | 11111044 |
| iMatrix-511 | Amsbio | 892 011 |
| LS Columns | Miltenyi Biotec | 130-042-401 |
| mTeSR plus | Stemcell Technologies | 100-0276 |

| | | |
|---|---|---|
| MTS assay | Abcam | ab197010 |
| Neural Crest Stem Cell MicroBeads | Miltenyi Biotec | 130-097-127 |
| Normal Goat Serum | Thermo Fisher | PCN5000 |
| Paraformaldehyde | Electron Microscopy Sciences | 15710 |
| ReLeSR | Stemcell Technologies | 5872 |
| Retinoic Acid | Sigma-Aldrich | R2625 |
| RIPA Lysis Buffer System | Santa Cruz Biotechnology | sc-24948 |
| Rock inhibitor (Y-27632) | R&D systems | 1254/10 |
| SB431542 | Tocris | 1614 |
| StemPro Accutase | Thermo Fisher Scientific | A1110501 |
| Triton X-100 | Millipore-Sigma | T8787 |
| Trypan blue | Thermo Fisher Scientific | T10282 |
| Tween 20 | Amresco | 0777 |

**Supplemental Table 2.** Primary antibodies used for immunoassays.

| Marker | Vendor | Host | Application | Dilution | Blocking |
|---|---|---|---|---|---|
| **OCT-3/4** | Santa Cruz sc-5279 | Mouse | ICC | 1:50 | +GS/+TR/+TW |
| **SSEA-4** | Abcam ab16287 | Mouse | ICC | 1:100 | +GS/−TR/−TW |
| **ZO-1** | ThermoFisher 40-2200 | Rabbit | ICC<br>WB | 1:100<br>1:50 | +GS/+TR/+TW<br>— |
| **NGFR** | Advanced Targeting Systems AB-N07 | Mouse | ICC | 1:1000 | −GS/−TR/−TW |
| **NG-2** | Millipore-Sigma MAB2029 | Mouse | ICC | 1:100 | +GS/+TR/+TW |
| **PDGFR-β** | Cell Signaling Technology 3169 | Rabbit | ICC | 1:100 | +GS/+TR/+TW |
| **γH2AX** | Abcam Ab11174 | Rabbit | WB | 1:50 | — |
| **NanoG** | R&D Systems AF1997 | Goat | WB | 1:50 | — |
| **Claudin-5** | Thermo Fisher 35-2500 | Mouse | WB | 1:50 | — |
| **GAPDH** | Abcam Ab181602 | Rabbit | WB | 1:1000 | — |

GS: normal goat serum; TR: Triton X-100; TW: Tween-20.

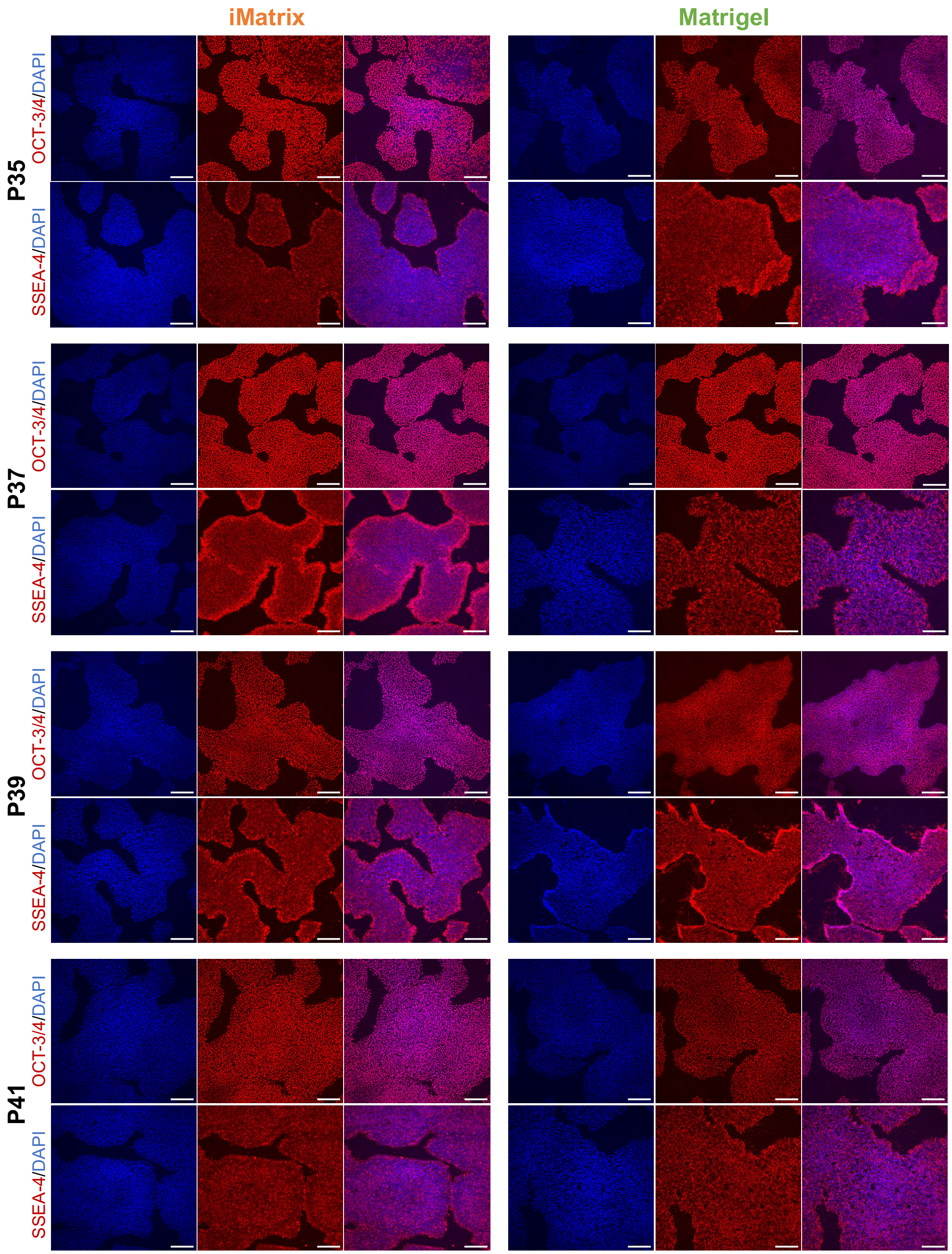

iMatrix
Matrigel
P35
OCT-3/4/DAPI
SSEA-4/DAPI
P37
OCT-3/4/DAPI
SSEA-4/DAPI
P39
OCT-3/4/DAPI
SSEA-4/DAPI
P41
OCT-3/4/DAPI
SSEA-4/DAPI

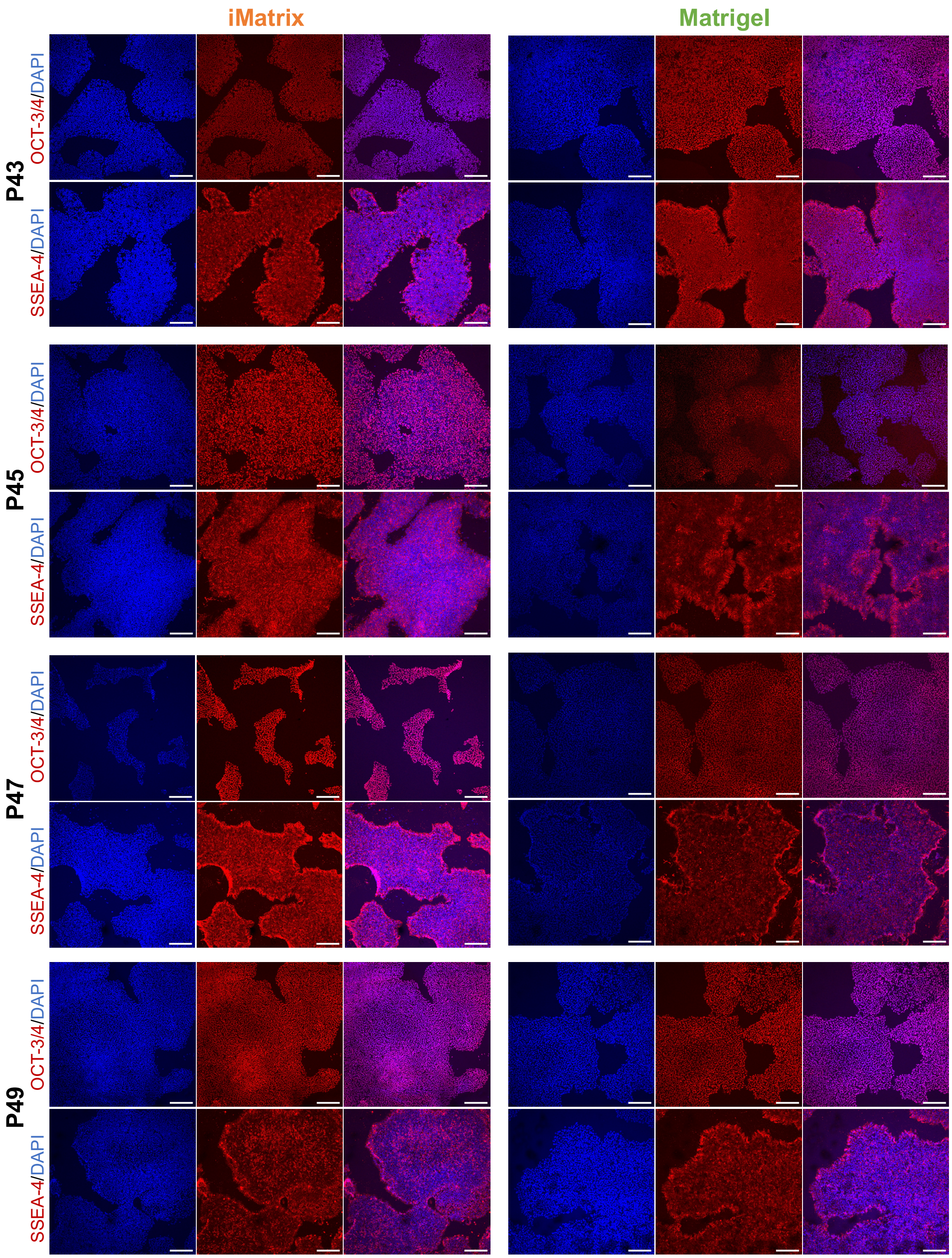


**Supplemental Figure 1.** Confocal microscopy images of iPSCs grown from passages 35 to 49 using iMatrix or Matrigel following immunoidentification of pluripotency markers SSEA-4 and OCT-3/4 (red fluorescence) and nuclear staining with DAPI (blue fluorescence). Scale bars: 200 µm. DAPI: 4',6-diamidino-2-phenylindole; OCT-3/4: octamer binding transcription factor-3/4; P: passage; SSEA-4: stage-specific embryonic antigen-4.

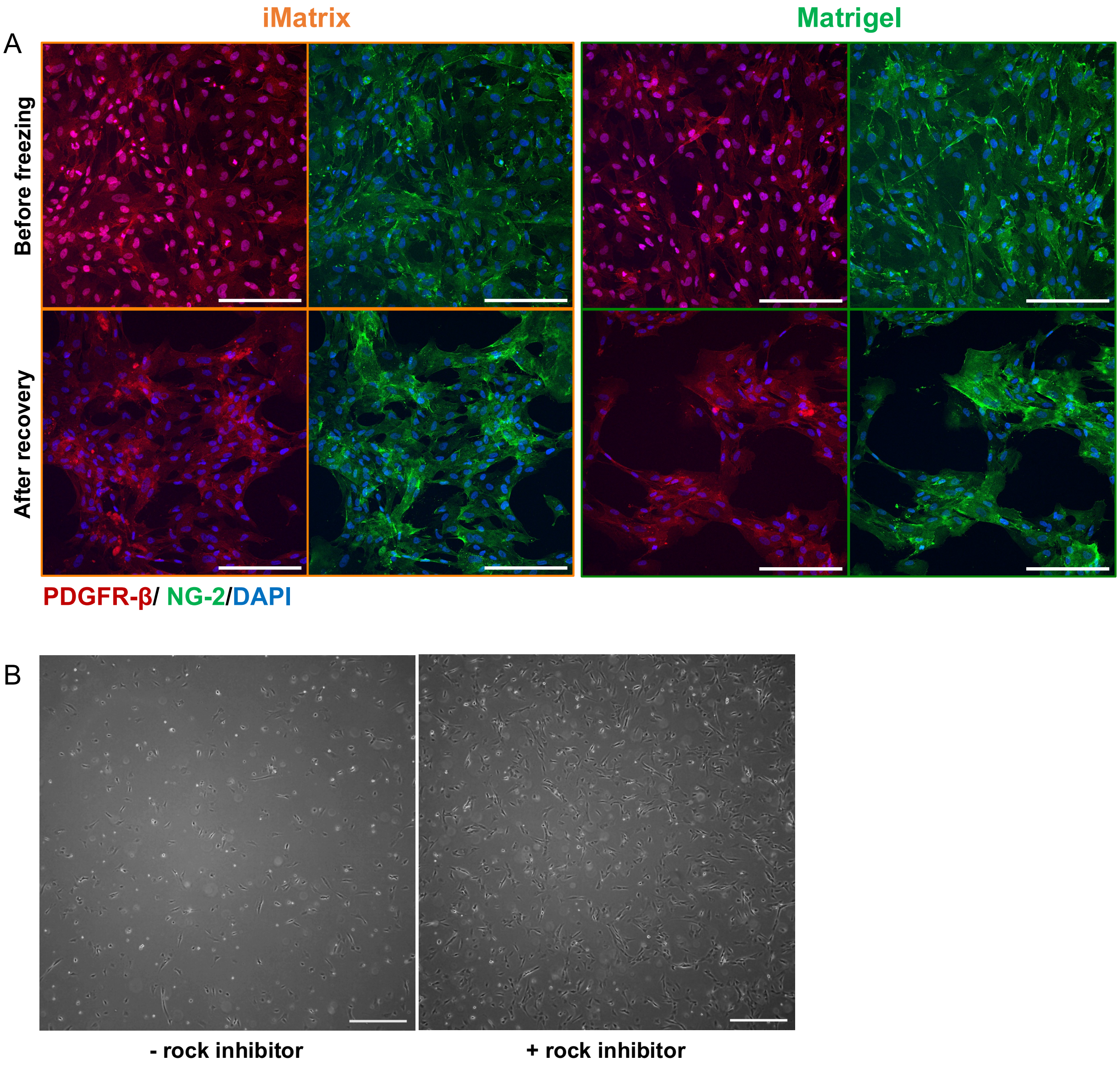


**Supplemental Figure 2. Cryopreservation of iPericytes**. **(A)** Confocal microscopy images of iM-iPericytes and MG-iPericytes before and after cryopreservation following immunolabeling of PDGFR-β (red fluorescence), NG-2 (green fluorescence), and nuclear staining (blue fluorescence). Scale bars: 200 µm. **(B)** Phase contrast images of iM-iPericytes frozen with or without rock inhibitor in the freezing media 48 hours after thawing. Scale bars: 500 µm. DAPI: 4',6-diamidino-2-phenylindole; NG-2: neural/glial antigen 2; PDGFR-β: platelet-derived growth factor receptor-β.
